# MANF stimulates autophagy and restores mitochondrial homeostasis to treat toxic proteinopathy

**DOI:** 10.1101/2023.01.10.523171

**Authors:** Yeawon Kim, Chuang Li, Chenjian Gu, Eric Tycksen, Anuradhika Puri, Terri A. Pietka, Jothilingam Sivapackiam, Yili Fang, Kendrah Kidd, Sun-Ji Park, Bryce G. Johnson, Stanislav Kmoch, Jeremy S. Duffield, Anthony J. Bleyer, Meredith E. Jackrel, Fumihiko Urano, Vijay Sharma, Maria Lindahl, Ying Maggie Chen

## Abstract

Misfolded protein aggregates may cause toxic proteinopathy, including autosomal dominant tubulointerstitial kidney disease due to uromodulin mutations (ADTKD-*UMOD*), one of the leading hereditary kidney diseases, and Alzheimer’s disease etc. There are no targeted therapies. ADTKD is also a genetic form of renal fibrosis and chronic kidney disease, which affects 500 million people worldwide. For the first time, in our newly generated mouse model recapitulating human ADTKD-*UMOD* carrying a leading *UMOD* deletion mutation, we show that autophagy/mitophagy and mitochondrial biogenesis are severely impaired, leading to cGAS-STING activation and tubular injury. Mesencephalic astrocyte-derived neurotrophic factor (MANF) is a novel endoplasmic reticulum stress-regulated secreted protein. We provide the first study that inducible tubular overexpression of MANF after the onset of disease stimulates autophagy/mitophagy and clearance of the misfolded UMOD, and promotes mitochondrial biogenesis through p-AMPK enhancement, resulting in protection of kidney function. Conversely, genetic ablation of endogenous MANF upregulated in the mutant mouse and human tubular cells worsens autophagy suppression and kidney fibrosis. Together, we discover MANF as a novel biotherapeutic protein and elucidate previously unknown mechanisms of MANF in regulating organelle homeostasis to treat ADTKD, which may have broad therapeutic application to treat various proteinopathies.

## INTRODUCTION

Misfolded protein due to genetic mutations and the resultant endoplasmic reticulum (ER) stress represent one important cause of ER storage disease and toxic proteinopathy, including autosomal dominant tubulointerstitial kidney disease due to uromodulin mutations (ADTKD- *UMOD*), Alzheimer’s disease ^1^, amyotrophic lateral sclerosis ^2^, Wolfram syndrome ^3^, cystic fibrosis ^4^, and α1-antitrypsin deficiency ^5^. Currently there is no mechanistic treatment. ADTKD- *UMOD* is a monogenic form of renal tubulointerstitial fibrosis and chronic kidney disease (CKD), which occurs in ∼10% of the population associated with significant morbidity and mortality ^6^. ADTKD also represents as many as 25% of patients with inherited kidney disease, after exclusion of polycystic kidney disease and Alport syndrome ^7^. Uromodulin (Tamm-Horsfall protein) is largely expressed in the thick ascending limb (TAL) tubular epithelial cells. Currently 135 *UMOD* mutations have been identified, and some of these mutations have been shown to cause protein misfolding and ER stress, eventually causing TAL damage, inflammatory cell infiltration and fibrosis ^8, 9^. However, the molecular link between the ER stress activation and renal fibrosis is still missing.

ER protein aggregates can activate the unfolded protein response (UPR). The UPR is regulated by three pathways: inositol-requiring enzyme 1 (IRE1)-spliced XBP1 (XBP1s), protein kinase-like ER kinase (PERK)-activating transcription factor 4 (ATF4), as well as cleavage of 90-kD activating transcription factor 6 (ATF6) to the active 50-kD ATF6. Meanwhile, there is intensive crosstalk between ER stress signaling and the autophagy-lysosomal pathway, and autophagy is a highly conserved protein degradation process responsible for removal of aggregate-prone proteins (aggrephagy) and damaged organelles. The autophagic flux proceeds through several phases: phagophore initiation and nucleation that requires unc-51-like kinase 1 and 2 (ULK1/2)/FIP200/Atg13 and PI3K/Vps34/Vps15/beclin 1, respectively; vesicle elongation forming autophagosomes that requires microtubule-associated proteins 1A/1B light chain 3B (LC3B) and Atg5-Atg12-Atg16 ubiquitin-conjugation systems; autophagosome maturation; and autophagosome-lysosome fusion forming autolysosomes ^10^. The mechanisms that regulate proteostasis of UMOD remain poorly understood and treatment is still lacking.

The selective removal of damaged mitochondria through autophagy is called mitophagy. Mitophagy is mainly mediated by LC3-associated, ubiquitin- and receptor-dependent pathways. PINK1, a mitochondrial Ser/Thr kinase, on the outer membrane of damaged mitochondria can recruit Parkin, an E3 ubiquitin ligase. After ubiquitination, damaged mitochondria are selectively recognized by adaptor proteins and engulfed by autophagosomes. Recently, it has been shown that mitofusin 2 (MFN2) is an additional PINK1 substrate for Parkin recruitment besides its role in mitochondrial fusion ^11^. Mitophagy receptors include BCL-2/adenovirus E1B 19 kDa protein-interacting protein 3 (BNIP3), BCL-2-interacting protein 3-like (BNIP3L) and FUN14 domain-containing 1 (FUNDC1) ^12^. Whether mitophagy is dysregulated in ADTKD-*UMOD* has not been studied.

Mesencephalic astrocyte-derived neurotrophic factor (MANF), an 18 kD, novel ER soluble protein, can exert protective function in Parkinson’s disease and ischemic stroke ^13, 14^, and regulate immune homeostasis in aging and retinal regenerative therapies ^15, 16^ in animal models. It is also upregulated and secreted in response to ER stress. The secreted extracellular MANF can protect cells from stress-induced cell death in experimental models of myocardial infarction ^17^ and diabetes ^18^ by inhibiting ER stress-induced apoptosis. We have discovered that secreted MANF can serve as a urinary ER stress biomarker ^19^ and function as an ER calcium stabilizer for ER-stressed podocytes *in vitro* ^20^. However, the function of MANF has not been investigated in kidney disease *in vivo*.

By using CRISPR/Cas9, we first generated an ADTKD-*UMOD* mouse model carrying UMOD p.Tyr178_Arg186 del, the mouse equivalent of the human UMOD p.His177_Arg185 del in-frame deletion, one of the most prevalent mutations in human ADTKD-*UMOD* ^21^. To assess the functional role of MANF in ADTKD, we generated both TAL cell-specific MANF knockout and inducible tubular cell-specific MANF transgenic mice. We demonstrate that MANF expression is increased exclusively in ER-stressed TAL epithelium. MANF deletion in mutant TALs exacerbates autophagy failure and renal fibrosis. Conversely, MANF overexpression in renal tubules augments autophagy/mitophagy, improves mitochondrial function, and inhibits tubular cGAS-STING dependent inflammation, thus attenuating kidney injury and fibrosis. For the first time, our findings reveal the role of MANF in maintaining functional autophagy, and highlight the important therapeutic function of MANF by regulating organelle homeostasis for the treatment of ADTKD and possibly other toxic proteinopathy.

## RESULTS

### Generation of UMOD Y178-R186 deletion mice to recapitulate human ADTKD-*UMOD*

We used CRISPR/Cas9 with one guide RNA (sgRNA) targeting the *Umod* gene’s 5’- TATGAGACCCTGACTGAGTACTGGCGC-3’ to delete amino acids YETLTEYWR in exon 3 (Figure 1A). The sgRNA and Cas9 protein that were complexed to generate the ribonucleoprotein, along with a 200-bp single-stranded donor DNA containing the Y178-R186 del, 27-bp deletion of *Umod*, were injected into C57BL/6J mouse embryos at the single-cell stage. Founder genotyping was performed by deep sequencing, and 32 founders were identified. The positive founders were further crossed to C57BL/6J mice to generate heterozygous F1 offspring, which were also deep sequenced to confirm correctly targeted alleles. The heterozygous mice of the F1 generation were found to have a germline transmission of 50%.

**Figure 1.**
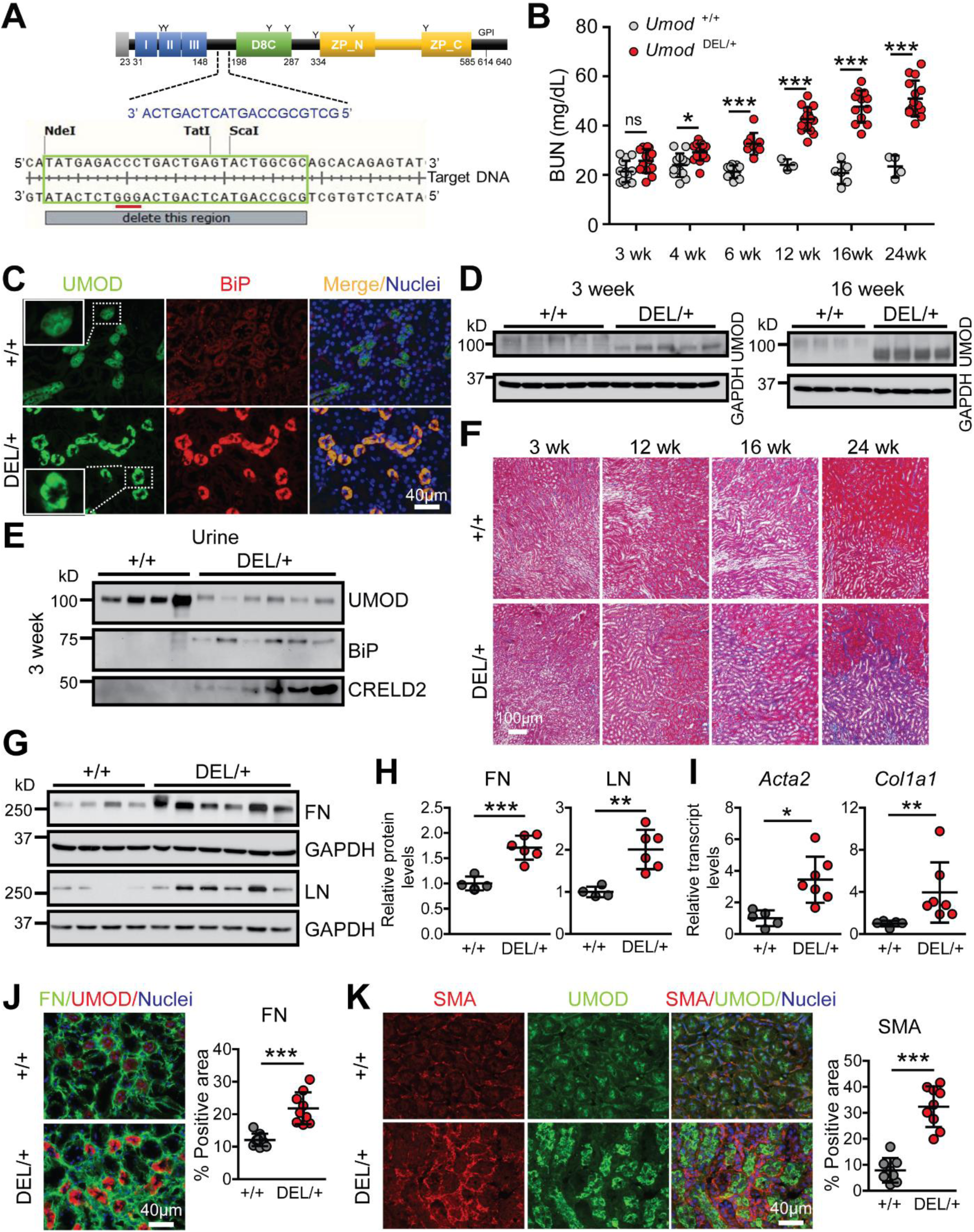
Generation of a mouse model that recapitulates human ADTKD-*UMOD*. (**A**) Schematic illustration of *Umod* genomic structure. The clustered regulatory interspaced short palindromic repeats (CRISPR) target sites are outlined in green, the sgRNA sequences are marked in blue, and protospacer adjacent motifs (PAMs) are underlined in red. Y indicates N-linked glycosylation sites. (**B**) BUN measurements over a 24-week period. Mean ± SD (n=4-17 mice/genotype). ns, not significant; *p<0.05; ***p<0.001. (**C**) Representative IF images of paraffin kidney sections stained for UMOD (green) and BiP (red) at 16 weeks. Scale bar, 40 µm. Insets magnify punctate, mutant UMOD staining. (**D**) Immunoblot to detect UMOD protein from whole-kidney tissues; the WT glycosylated band migrated slower than the mutant under-glycosylated band. n=4-5 mice/genotype. (**E**) WBs showed excretion of urinary UMOD, BiP and CRELD2 from *Umod ^+/+^* and *Umod* ^DEL/+^ mice at 3 weeks of age. The urinary excretion of these ER stress biomarkers was normalized to urine creatinine excretion such that the urine volume applied to the gel reflected the amount of urine containing 4 µg of creatinine. n=4-6 mice/genotype. (**F**) Representative Masson’s trichrome staining of paraffin kidney sections from *Umod ^+/+^* and *Umod* ^DEL/+^ mice from 3 to 24 weeks. Scale bar, 100 µm. (**G-H**) WBs of whole-kidney lysates from *Umod ^+/+^* and *Umod* ^DEL/+^ mice at 16 weeks to detect FN and LN (**G**) with densitometry analysis (**H**). The average FN or LN/β-actin ratio in WT mice was set as 1. Mean ± SD (n=4-6 mice/genotype). **p<0.01; ***p<0.001. (**I**) Quantitative RT-PCR analysis of relative transcript levels of *Acta2* and *Col1a1* in *Umod ^+/+^* and *Umod* ^DEL/+^ kidneys at 16 weeks of age. Gene expression was normalized to 18s. Mean ± SD (n=5-7 mice/genotype). *p<0.05; **p<0.01. (**J**) Dual IF staining of FN (green) and UMOD (red) with a nuclear counterstain (Hoechst 33342, blue) on frozen kidney sections from *Umod ^+/+^* and *Umod* ^DEL/+^ mice at 16 weeks with quantification. Scale bar, 40 µm. Mean ± SD (n=10 images/genotype). ***p<0.001. (**K**) Dual IF staining of SMA (red) and UMOD (green) with nuclear counterstain (blue) on frozen kidney sections from *Umod ^+/+^* and *Umod* ^DEL/+^ mice at 24 weeks with quantification. Scale bar, 40 µm. Mean ± SD (n=10 images/genotype). ***p<0.001.

A cohort of heterozygous (*Umod* ^DEL/+^) and WT littermates were followed until 24 weeks of age. Blood urea nitrogen (BUN) was slightly increased at 3 weeks, significantly elevated by 4 weeks, and progressively increased between 4 and 24 weeks in the mutants (Figure 1B). As expected, co-immunofluorescence (IF) staining of UMOD and the ER stress marker BiP in kidney sections at 16 weeks highlighted discontinuous and punctate UMOD protein aggregates in the ER in the mutant mice (Figure 1C, insets), consistent with our previous finding from a human kidney biopsy carrying the *UMOD* H177-R185 del ^19^. In addition, the mutant UMOD induced ER stress (Figure 1C). Meanwhile, we also noted that the mutant UMOD aggregation occurred by 3 weeks (Supplemental Figure 1A). In agreement with the staining result, WB revealed elevation of the under-glycosylated form of mutant UMOD due to ER retention as early as 3 weeks and much more prominent ER accumulation at 16 weeks (Figure 1D). In contrast, urinary UMOD excretion was markedly decreased in the mutants, along with increased excretion of our recently identified urinary ER stress biomarkers, secreted BiP ^22^ and cysteine-rich with EGF-like domains 2 (CRELD2) ^23^ as early as 3 weeks (Figure 1E). Masson’s trichrome staining showed progressive tubulointerstitial fibrosis during the disease progression, with the fibrosis accentuated in the corticomedullary junction (Figure 1F). Consistent with this observation, whole-kidney mRNA and protein quantifications and IF staining demonstrated a progressive increase in fibrosis markers in DEL/+ mice. By 12 weeks, fibronectin (FN) expression was increased (Supplemental Figure 1B). At 16 weeks, a marked increase in both protein levels of FN and laminin (LN) (Figure 1, G, H and J) and transcript levels of smooth muscle actin (SMA) (*Acta2*) and collagen I α1 (*Col1a1*) (Figure 1I) were observed. At 24 weeks, immunoblots of LN, SMA and FN (Supplemental Figure 1C), q-PCR of *Col1a1* and *Tgfb* (Supplemental Figure 1D) and IF of SMA (Figure 1K) analyses continue to display the substantial increase in fibrosis in DEL/+ mice. Overall, we have successfully established an ADTKD-*UMOD* mouse model harboring a pathogenic human mutation that recapitulates monogenic CKD in the patients.

### Impaired autophagy in TALs expressing the mutant UMOD in ADTKD-*UMOD* mice

To investigate the molecular mechanisms underlying the disease pathogenesis, first, we determined which UPR branch was activated. WB demonstrated that the ATF6 pathway was activated by 12 weeks (Figure 2A, arrow), and persisted through 24 weeks in the mutant kidneys, whereas the other two UPR branches were not involved (Supplemental Figure 2, A-C). Next, we examined autophagic activity in the mutant kidneys. As elevated LC3B-II levels, which indicate autophagosomal accumulation, can signify the changes in both formation and degradation of the autophagosomes, p62, the selective degradation substrate of autophagy that links ubiquitinated proteins to LC3, also needs to be assessed. Accumulation of p62 protein in the absence of increased transcription indicates that autophagy clearance is inhibited. During the disease progression, protein abundance of LC3-II and p62 was increased by 12 weeks (Supplemental Figure 2D) and persisted through 16-24 weeks (Figure 2B and Supplemental Figure 2E) without transcriptional upregulation of P62 (Supplemental Figure 2F) in the mutant kidneys, demonstrating impaired autophagic degradation of the misfolded protein.

**Figure 2.**
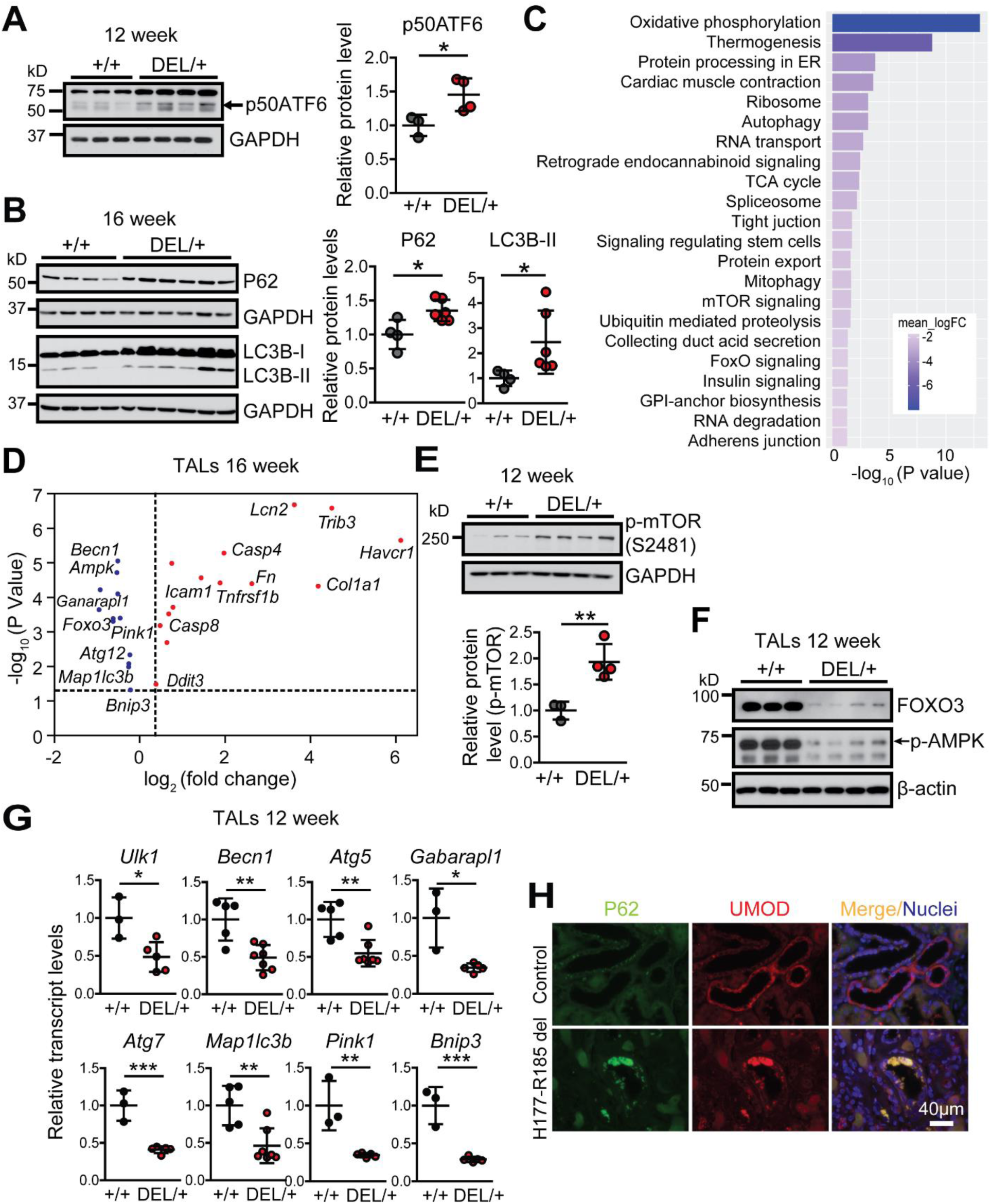
Impaired autophagy in the mutant TALs in ADTKD-*UMOD*. (**A**) Whole-kidney lysates from *Umod ^+/+^* and *Umod* ^DEL/+^ mice at 12 weeks were analyzed by WB for active, cleaved p50ATF6 (arrow) with densitometry analysis. The average p50ATF6/β-actin ratio in WT mice was set as 1. Mean ± SD (n=3-4 mice/genotype). *p<0.05. (**B**) WBs of whole-kidney lysates at 16 weeks to detect the autophagy mediators P62 and LC3B (LC3B-I inactive, LC3B-II active) with densitometry analysis. The average p62 or LC3B-II/β-actin ratio in WT mice was set as 1. Mean ± SD (n=4-6 mice/genotype). *p<0.05. (**C-D**) RNA-seq analysis of mutant and WT UMOD-expressing TAL epithelium at 16 weeks. n=5 per group. (**C**) KEGG pathway analysis. The top 22 downregulated pathways were obtained from the KEGG analysis of the log2 fold-changes in primary TAL cells isolated from *Umod* ^DEL/+^ and *Umod* ^+/+^ mice at 16 weeks. (**D**) RNA-seq analysis of genes for autophagy, inflammation, ER stress response and fibrosis in the mutant TALs compared with WT TALs (Benjamini-Hochberg adjusted p value<0.05). (**E**) WB from *Umod ^+/+^* and *Umod* ^DEL/+^ kidneys for levels of p-mTOR (S2481) at 12 weeks with densitometry analysis. The average p-mTOR/β-actin ratio in WT mice was set as 1. Mean ± SD (n=3-4 mice/genotype). **p<0.01. (**F**) WBs from *Umod ^+/+^* and *Umod* ^DEL/+^ primary TALs for levels of FOXO3 and p-AMPK (arrow) at 12 weeks. n=3-4 mice/genotype. (**G**) Transcript analysis of a panel of autophagy and mitophagy-related genes from isolated primary TAL cells at 12 weeks. Gene expression was normalized to 18s. Mean ± SD (n=3-7 mice/genotype). *p<0.05; **p<0.01; ***p<0.001. (**H**) Representative IF images of human kidney biopsies obtained from a patient with p.H177- R185 del and from a normal kidney, stained for P62 (green) and UMOD (red) with nuclei counterstain (blue) on paraffin kidney sections. Scale bar, 40 µm.

To gain further insight into the mechanisms underlying the defective autophagic activity and its functional impact on the pathogenesis of disease, we performed RNA sequencing (RNA-seq) of mRNA isolated from UMOD-producing TAL cells purified from *Umod* ^DEL/+^ and *Umod* ^+/+^ littermates at 16 weeks, the full-blown stage of the disease. Principal components analysis of RNA-seq expression patterns across these two groups revealed clustering within each genotype, suggesting two very distinct cell populations (Supplemental Figure 3A). In addition, our analysis revealed that 9566 genes were differentially expressed between mutant and WT TAL cells (Benjamini-Hochberg adjusted p≤0.05) (Supplemental Figure 3B). As depicted in the volcano plot, among these 9566 differentially expressed transcripts, including 4706 upregulated and 4860 downregulated, 244 genes were up-regulated and 149 were down-regulated by 4-fold or more in the mutant UMOD-expressing TAL cells (Supplemental Figure 3B). Kyoto Encyclopedia of Genes and Genomes (KEGG) pathway analysis for primary mutant versus WT TAL cells revealed top 22 downregulated pathways, which are related to mitochondria oxidative phosphorylation, ATP generation (thermogenesis), ER proteostasis, autophagy, TCA cycle, mitophagy, mTOR and FOXO signaling, as well as ubiquitin-mediated protein degradation (Figure 2C). Additional RNA-seq analysis of dysregulated genes specifically in mutant TALs at the 16-week time point confirmed a significant suppression of autophagy-related genes, including *Becn1*, *Ampk*, *Gabarapl1* encoding ATG8, *Foxo3*, *Pink1*, *Atg12*, *Map1lc3b* encoding LC3B, and *Bnip3* (Figure 2D). We also observed a marked activation of genes involved in epithelial injury (*Lcn2* encoding neutrophil gelatinase-associated lipocalin (NGAL), *Havcr1* encoding Kim1), matrix dysregulation and fibrosis (*Col1a1*, *Fn*), inflammation (*Tnfrsf1b*, *Icam1*), ER stress-induced apoptosis (*Trib3*, *Ddit3*), as well as *Casp4* and *Casp8* (Figure 2D). Q- PCR analysis of the isolated TAL cells at 16 weeks confirmed repression of autophagy in the mutant TALs (Supplemental Figure 4).

mTOR negatively regulates autophagy through phosphorylation at Ser757 to inhibit ULK1 activity and autophagosome formation. Consistent with the RNA-seq data (Figure 2C), WB showed increased levels of active p-mTOR (S2481) in the mutant kidneys at 12 weeks, the early stage of the disease (Figure 2E). To further understand the molecular control of the dysfunctional autophagy in the mutant TALs, we purified primary WT and mutant TAL cells at early stage of the disease, 12 weeks. Given that in the mutant TALs RNA-seq revealed inhibited FOXO signaling (Figure 2C), a family of critical transcriptional factors that induce autophagy by transactivating expression of genes encoding induction, nucleation, elongation and fusion of the autophagic process ^24^, we examined expression of FOXO3a. The mutant TAL cells exhibited remarkably decreased FOXO3 expression at 12 weeks (Figure 2F). Moreover, protein expression of the active, phosphorylated AMP-activated protein kinase (p-AMPK) at Thr-172, which can phosphorylate FOXO3 at Ser413 or Ser588, promoting nuclear accumulation and stabilization and preventing cytoplasmic degradation of FOXO3 ^24^, as well as block mTOR activity, was also dramatically reduced in the mutant TAL cells at 12 weeks (Figure 2F, arrow). Besides the upstream regulator, a further transcriptional analysis of FOXO3 downstream targets in the mutant UMOD epithelium at 12 weeks revealed inhibition of a panel of autophagy machinery genes related to initiation (*Ulk1*), nucleation (*Becn1*) and elongation (*Atg5*, *Gabarapl1*, *Atg7* and *Map1lc3b*) of autophagosomes, as well as suppression of *Pink1* and *Bnip3*, which mediates mitophagy ^25^ (Figure 2G). Finally, we confirmed our results obtained from the mouse model in human kidney biopsies. Co-IF staining of P62 and UMOD clearly showed increased P62 aggregates in the UMOD-positive epithelium in a patient harboring H177-R185 del compared with a healthy control (Figure 2H). Taken together, these findings demonstrate that autophagy is actively suppressed in the UMOD proteinopathy model.

### Defective mitophagy and mitochondrial biogenesis in the mutant TALs

Mitophagy depends on the activity of the autophagy machine, and p-AMPK/FOXO3 regulate mitophagy as well ^26^, we therefore set out to study whether mitophagy is inhibited in the mutant TALs. Heatmap of the 16-week RNA-seq data showed that expression of multiple genes involved in mitophagy was significantly downregulated in the mutant vs. WT TALs (Benjamini-Hochberg adjusted p values≤0.05) (Figure 3A). Immunoblot analysis of isolated TALs at 12 weeks further validated that both ubiquitin-mediated PINK1/MFN2/Parkin pathway and receptor-mediated BNIP3/BNIP3L/FUNDC1 pathways were markedly inhibited in the mutant TALs (Figure 3B). Moreover, Mitochondrial fission is required for mitophagy and allows damaged mitochondria to be eliminated by mitophagy ^27^. In agreement with suppressed mitophagy, we also observed decreased expression of mitochondrial fission protein 1 (FIS1) in the mutant TALs at the early stage of disease, as evidenced by the heatmap (Figure 3A) and WB (Figure 3C).

**Figure 3.**
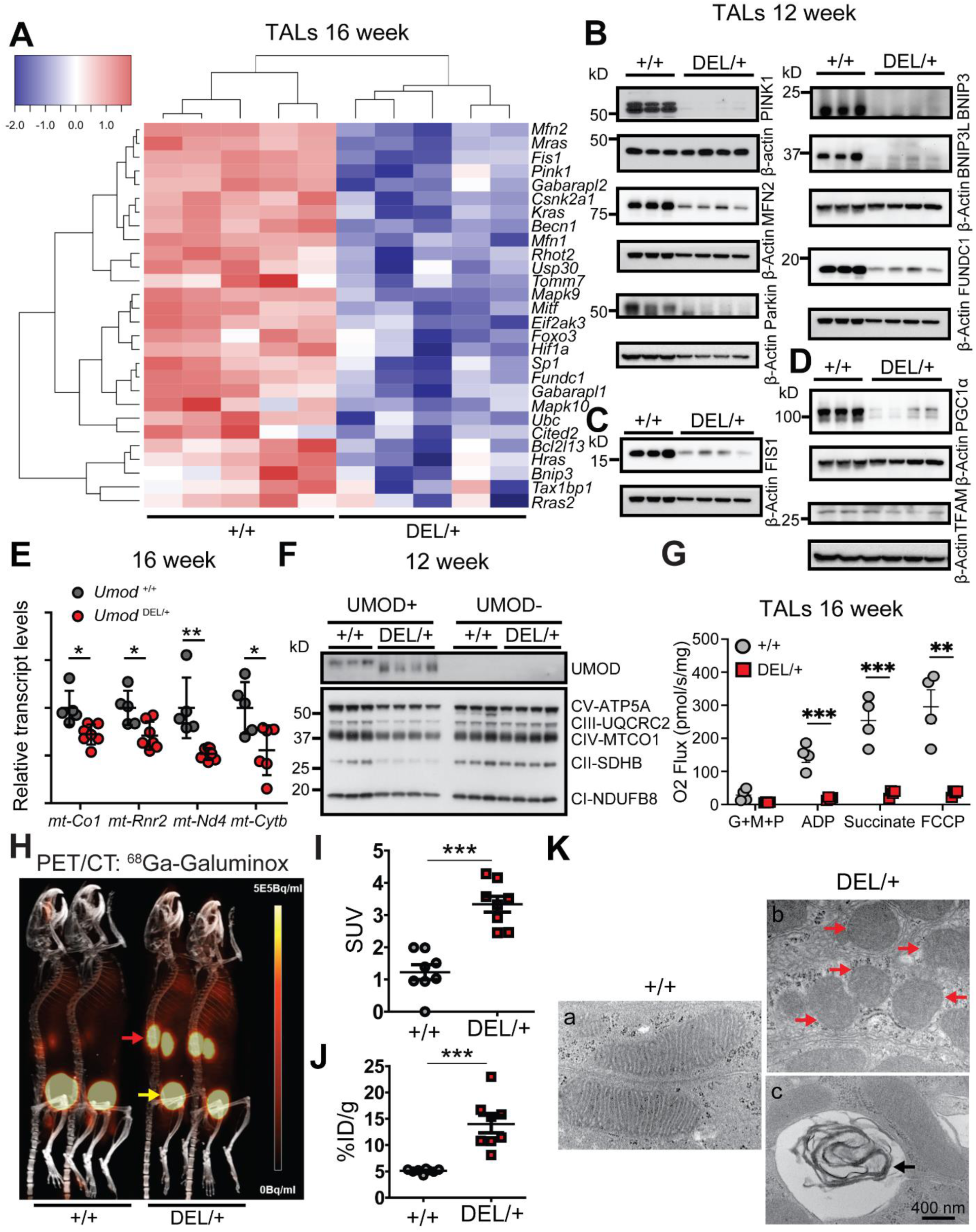
Deficient mitophagy and dysfunctional mitochondrial biogenesis in the mutant TALs in ADTKD. (**A**) Heatmap of downregulated mitophagy-associated genes with Benjamini-Hochberg adjusted p values≤0.05. RNA was isolated from *Umod ^+/+^* and *Umod* ^DEL/+^ TALs from 16- week-old mice. n=5 mice per group. (**B**) Representative immunoblot analysis monitoring mitophagy pathways, including PINK1, MFN2, Parkin, BNIP3, BNIP3L and FUNDC1 in *Umod ^+/+^* and *Umod* ^DEL/+^ TALs at 12 weeks. n=3-4 mice/genotype. (**C**) Representative immunoblot analysis monitoring mitochondrial fission marker FIS1 in *Umod ^+/+^* and *Umod* ^DEL/+^ TALs at 12 weeks. n=3-4 mice/genotype. (**D**) Representative immunoblot analysis monitoring mitochondrial biogenesis markers, PGC1α and TFAM in *Umod ^+/+^* and *Umod* ^DEL/+^ TALs at 12 weeks. n=3-4 mice/genotype. (**E**) Relative mRNA levels of mtDNA genes, *mt-Co1*, *mt-Rnr2*, *mt-Nd4* and *mt-Cytb* from *Umod ^+/+^* and *Umod* ^DEL/+^ kidneys at 16 weeks. Gene expression was normalized to 18s. Mean ± SD (n=5-7 mice/genotype). *p<0.05; **p<0.01. (**F**) Representative immunoblot analysis monitoring expression of mitochondrial respiratory complexes I-V by utilizing a total OXPHOS WB antibody cocktail in UMOD-positive and UMOD-negative renal tubular epithelium from *Umod ^+/+^* and *Umod* ^DEL/+^ kidneys at 12 weeks. n=3-4 mice/genotype. (**G**) Measurement of mitochondrial respiration using an OROBOROS Oxygraph system in permeabilized TALs from *Umod ^+/+^* and *Umod* ^DEL/+^ mice at 16 week following sequential additions of glutamate, malate and pyruvate (G+M+P); adenosine diphosphate (ADP); succinate and FCCP. Mean ± SD (n=4 mice/genotype). **p<0.01, ***p<0.001. (**H**) ^68^Ga-Galuminox (100 µCi) was injected via tail-vein into *Umod ^+/+^* and *Umod* ^DEL/+^ mice at 16 weeks. Static PET scans were acquired from 30-50 minutes post tail-vein injection. PET/CT images as shown are a summation of frames from 30-45 minutes. N=8 kidneys per group. Uptake of radiotracers in the kidneys (red arrows) and bladder (yellow arrows) were shown. (**I**) SUV analysis of ^68^Ga-Galuminox uptake in kidneys of the indicated genotypes. Mean ± SD (n = 8 kidneys/genotype). ***P<0.001. (**J**) Post-PET imaging biodistribution data (%ID/g) of *Umod ^+/+^* and *Umod* ^DEL/+^ kidneys at 16 weeks. Mean ± SD (n = 8 kidneys/genotype). ***P<0.001. (**K**) TEM ultrastructural analysis in the renal tubules from *Umod ^+/+^* (**a**) and *Umod* ^DEL/+^ **(b-c)** mice at 16 weeks. Red arrows indicate aggregates of swollen mitochondria with disrupted cristae (**b**), and black arrow indicates “myelin body” (**c**). Scale bar, 400 nm.

Given that during stress, AMPK also activates peroxisome proliferator-activated receptor-γ (PPARγ) co-activator 1α (PGC1α), which regulates mitochondrial biogenesis genes through interaction with PPARγ or estrogen-related receptors (ERRs) ^26^, we next examined mitochondrial biogenesis in our ADTKD-*UMOD* mouse model. At 12 weeks, both PGC1α and its downstream target mitochondrial transcription factor A (TFAM), a key regulator of mitochondrial gene expression ^28^, were substantially repressed in the mutant TALs compared with WT TALs (Figure 3D). Consequently, transcript levels of mitochondrial DNA (mtDNA), such as *mt-Co1*, *mt-Rnr2*, *mt-Nd4* and *mt-Cytb* in the mutant kidneys at 16 weeks (Figure 3E), as well as protein levels of mitochondrial oxidative phosphorylation (OXPHOS)-related proteins in the mutant TALs at 12 weeks (Figure 3F) were significantly lower than controls. In contrast, in UMOD-negative tubular cells, no difference in expression of the subunits of electron transport complex (ETC) proteins was noted between WT and mutant kidneys (Figure 3F). In line with the WB result, the KEGG analysis clearly depicted downregulation of all five ETCs in the mutant TALs as compared to WT TALs at 16 weeks (Supplemental Figure 5).

To determine the activity of ETCs and mitochondrial respiratory function, we performed high-resolution respirometry using an OROBOROS Instruments Oxygraph-O_2_k system in freshly isolated WT and mutant TAL cells at 16 weeks. To measure O_2_ flux, different substrates were added sequentially, including glutamate, malate and pyruvate (G+M+P), ADP, and succinate, followed by the OXPHOS uncoupler carbonyl cyanide-p-trifluoromethoxyphenylhydrazone (FCCP) to determine leak respiration, complex I activity, complex I+II activity and maximal ETC capacity, respectively. As shown in Figure 3G, oxygen consumption after addition of ADP and succinate substrates, as well as FCCP was significantly decreased in the mutant TALs compared to that in WT TALs, indicating disruption of mitochondrial respiratory function in the mutant TALs (Figure 3G). By contrast, there was no difference in oxygen consumption for UMOD-negative epithelial cells between WT and mutant kidneys at 16 weeks (Supplemental Figure 6).

Mitophagy blockade can lead to accumulation of damaged, reactive oxygen species (ROS)- generating mitochondria. Meanwhile, lack of efficient oxidative phosphorylation due to impaired mitochondrial biogenesis can result in overproduction of mitochondrial ROS. Thus, we anticipated increased accumulation of mitochondrial ROS in the mutant kidneys. To prove our hypothesis unambiguously, we employed noninvasive, sensitive, and quantitative PET/CT molecular imaging to detect mitochondrial ROS by utilizing our newly developed mitochondrial ROS radiotracer ^68^Ga-Galuminox ^29, 30^. To our knowledge, this is the first ^68^Ga-radiotracer (incorporated with a nonconventional, generator-produced isotope) capable of detecting mitochondrial ROS in live animals. Representative preclinical PET/CT images (summation of frames over 30-45 minutes) post-administration of ^68^Ga-Galuminox were shown in Figure 3H (red arrow). ^68^Ga-Galuminox showed a 2.73 fold higher uptake in kidneys of *Umod* ^DEL/+^ mice (standard uptake value (SUV): 3.33 ± 0.25, n=8) compared with WT littermates (SUV: 1.22 ± 0.23, n=8, *P*<0.0001) (Figure 3I). For further correlating preclinical PET imaging data, post-imaging biodistribution studies were also conducted and percentage of activity remained in kidneys at 1 hour post injection of the radiotracer was quantified as the percentage injected dose (% ID) per gram of kidney. ^68^Ga-Galuminox demonstrated that compared with the WT kidneys (% ID/g: 5.1 ± 0.14, n=8), the radiotracer was retained 2.7-fold higher in the mutant kidneys (% ID/g: 13.99 ± 1.66, n=8, *P*<0.001) (Figure 3J).

Consistent with these biochemical, functional and *in vivo* molecular imaging studies, transmission electron microscopy (TEM) ultrastructural analysis of mitochondrial integrity showed that mitochondria were aligned in well-preserved rows in normal renal tubules (Figure 3Ka). Mutant kidneys at 16 weeks showed disorganized mitochondrial arrays and aggregates of swollen mitochondria with disrupted cristae (Figure 3Kb, red arrows), as well as “myelin body” suggesting possible lysosome damage (Figure 3Kc, black arrow). Collectively, these data indicate impaired mitophagy and mitochondrial biogenesis/OXPHOS, leading to increased level of mitochondrial ROS and aberrant mitochondrial ultrastructure in the mutant TALs.

### Increased inflammation and activation of cGAS-STING signaling in ADTKD-*UMOD* mice

Gene Ontology (GO) analysis of the log2 fold-changes for primary mutant versus WT TAL cells at 16 weeks demonstrated that the most upregulated pathways in the mutant TAL cells were related to immune response and extracellular matrix organization (Figure 4A). Q-PCR analysis of the isolated TAL cells at 16 weeks confirmed activation of inflammatory genes, such as tumor necrosis factor (TNFα), interleukin-1β (IL-1β), interleukin-6 (IL-6), intercellular adhesion molecule 1 (ICAM1) and chemokine (C-C motif) ligand 2 (CCL2) in the mutant TALs (Figure 4B). Consistent with increased transcript levels of inflammatory cytokines, PAS staining (Figure 4C, red arrow) at 24 weeks highlighted kidney ingress of massive inflammatory cells and few kidney cyst (Figure 4C, black arrow), reminiscent of findings noted in human kidney biopsies ^23^. Moreover, co-IF staining of the macrophage marker F4/80 with UMOD showed macrophage infiltration surrounding mutant TAL tubules at 24 weeks (Figure 4, D-E).

**Figure 4.**
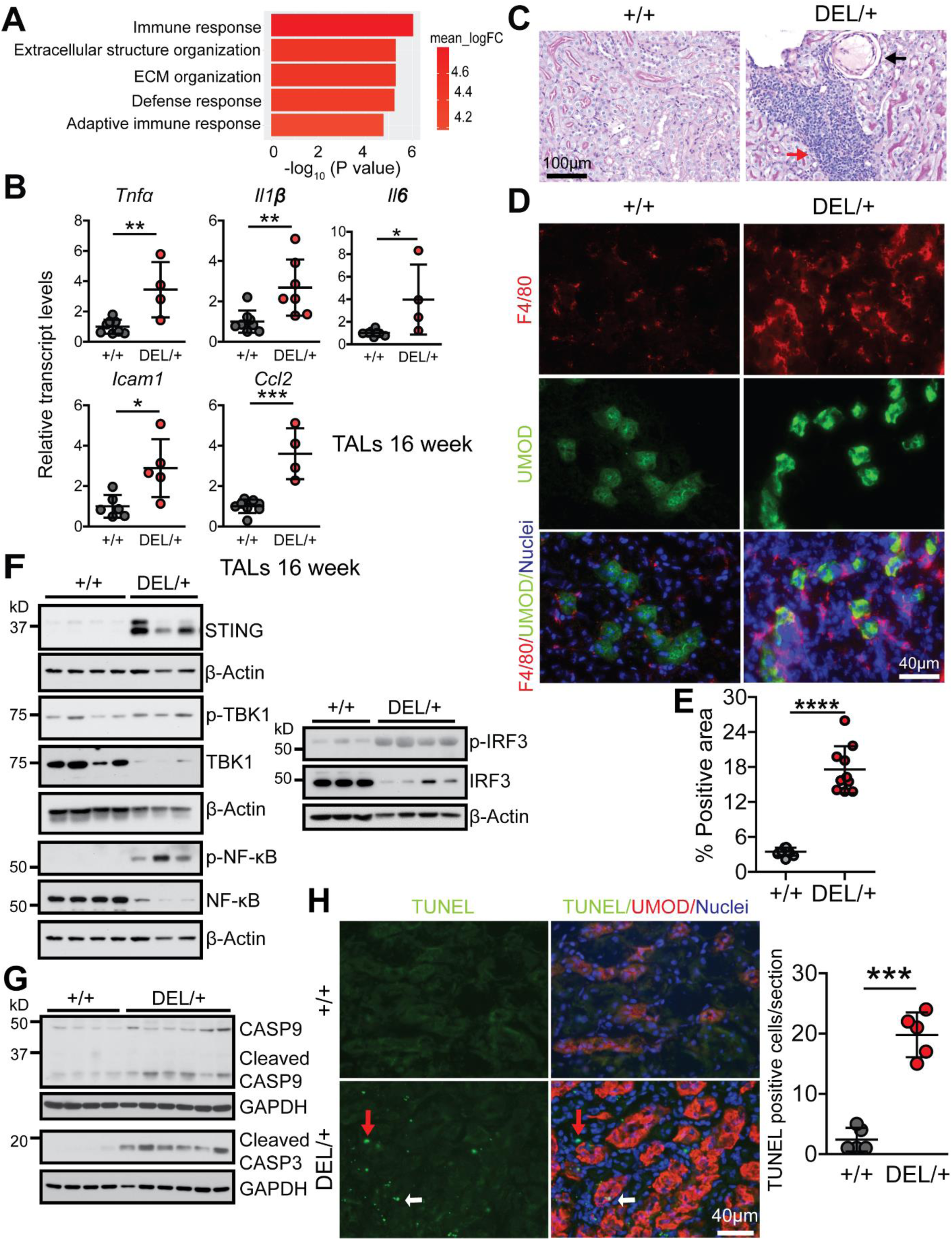
Activation of STING signaling, increased inflammation and apoptosis in ADTKD-*UMOD*. **(A)** RNA-seq analysis of WT and mutant TAL epithelium at 16 weeks. n=5 per group. GO biological process perturbation bar plot showed the most upregulated pathways in the mutant vs. WT TALs at 16 weeks. **(B)** Quantitative PCR for proinflammatory cytokines of isolated primary TAL cells at 16 weeks. Gene expression was normalized to β-actin. Mean ± SD (n=4-8 mice/genotype). *p<0.05; **p< 0.01; ***p<0.001. **(C)** PAS staining of paraffin kidney sections from *Umod ^+/+^* and *Umod* ^DEL/+^ mice at 24 weeks. Red arrow indicates interstitial inflammatory infiltration. Black arrow indicates a cortical renal cyst. Scale bar, 100 μm. **(D-E)** Representative IF staining of macrophage (F4/80, red) and UMOD (green) with nuclei counterstain (blue) on frozen kidney sections from *Umod ^+/+^* and *Umod* ^DEL/+^ mice at 24 weeks **(D)** with quantification **(E)**. Scale bar, 40 µm. Mean ± SD (n=10 images/genotype). ****p<0.0001. **(F)** Immunoblot analysis monitoring STING pathway and its downstream targets, including STING, p-TBK1/TBK1, p-NF-κB p65/NF-κB p65 and p-IRF3/IRF3 in *Umod ^+/+^* and *Umod* ^DEL/+^ TALs at 16 weeks. **_(G)_** Immunoblot analysis monitoring cleaved CASP9 and CASP3 in *Umod ^+/+^* and *Umod* ^DEL/+^ kidneys at 16 weeks. n=4-6 mice/genotype. (**H**)Representative IF staining of TUNEL (green) and UMOD (red) with nuclei counterstain (blue) on frozen kidney sections from *Umod ^+/+^* and *Umod* ^DEL/+^ mice at 24 weeks. TUNEL staining easily detected increased apoptosis in both UMOD^+^ (white arrows) and UMOD^-^ cells (red arrows) at 24 weeks. Scale bar, 40 µm. The TUNEL-positive cells were quantified. Mean ± SD (n=5 images/genotype). ***p<0.001.

To explore the molecular mechanism underpinning the increased immune response in the mutant kidneys, we examined cGAS (cyclic guanosine monophosphate-adenosine monophosphate synthase)-stimulator of interferon genes (STING) signaling that may be activated following the failure of mitochondrial quality control (Figure 3). Cytosolic mtDNA can be sensed by cGAS, which induces synthesis of cyclic guanosine adenosine monophosphate (cGAMP). cGAMP binds to ER adapter protein STING, and activated STING migrates from ER to the Golgi apparatus. During this process, STING can recruit and activate TANK-binding kinase 1 (TBK1), which in turn, activates the downstream NF-κB and interferon (IFN) regulatory factor 3 (IRF3) signal cascades, thus inducing the expression of inflammatory factors and type I IFN to strengthen immune responses ^31^. Indeed, STING, pTBK1/TBK1 ratio and their downstream targets p-NF-κB p65 and p-IRF3 exhibited increased abundance in the mutant TALs compared with WT TALs at 16 weeks (Figure 4F). It has been shown that disruption of mitochondrial integrity and activation of STING and inflammation in renal tubules strongly contribute to cell death and kidney failure ^32, 33^. Similarly, we observed a prominent cleavage of procaspase-9 in the mutant kidneys at 16 weeks, indicating activation of the mitochondria-dependent apoptotic pathway (Figure 4G). Ultimately, the executioner caspase-3 was activated at 16 weeks (Figure 4G), and TUNEL staining easily detected increased apoptosis in both UMOD^+^ (Figure 4H, white arrows) and UMOD^-^ cells (Figure 4H, red arrows) at 24 weeks. In summary, a defective engagement of mitophagy and mitochondrial biogenesis in response to ER stress triggered by the misfolded UMOD stimulates cGAS-STING signaling and innate immune response, which may eventually lead to tubular cell death and fibrosis in ADTKD-*UMOD*.

### mtDNA leakage into the cytosol activates cGAS-STING in ADTKD-*UMOD*

To directly demonstrate that STING pathway is activated by cytosolic leakage of mtDNA and recapitulate the *in vivo* findings in our mouse model, we generated a cellular model of ADTKD- *UMOD* in HEK 293 cells by transducing the cells with lentivirus expressing WT or mutant UMOD (H177-R185 del) fused to N-terminal GFP. It has been shown previously that primary TAL cells lose UMOD expression after 10 days of *in-vitro* culturing ^9^, and that immortalized mouse TAL cell line does not express UMOD ^34^. Thus, *UMOD* expression was placed under the control of the cytomegalovirus (CMV) promoter and secretion of the GFP-UMOD fusion proteins was directed by the endogenous secretory signal peptide of UMOD. Confocal live imaging of stable cell lines showed clear membrane enrichment for WT UMOD and diffuse reticular distribution in the cytoplasm for the mutant UMOD, whereas GFP alone was located in both cytoplasm and nucleus (Figure 5A). Q-PCR demonstrated that the mRNA expression level of *UMOD* in the two cell lines was comparable (Figure 5B). Immunoblot analysis showed elevation of mutant forms of both UMOD and GFP in the mutant H177-R185 del UMOD cell line (DEL) compared to WT cell line, associated with reduced secretion of the GFP-tagged mutant UMOD into the cell culture medium (Figure 5C). IF staining following fixation and permeabilization of the stably transduced HEK 293 cells confirmed ER retention of the mutant UMOD by its increased co-localization with the ER marker calnexin as compared to WT UMOD (Figure 5D).

**Figure 5.**
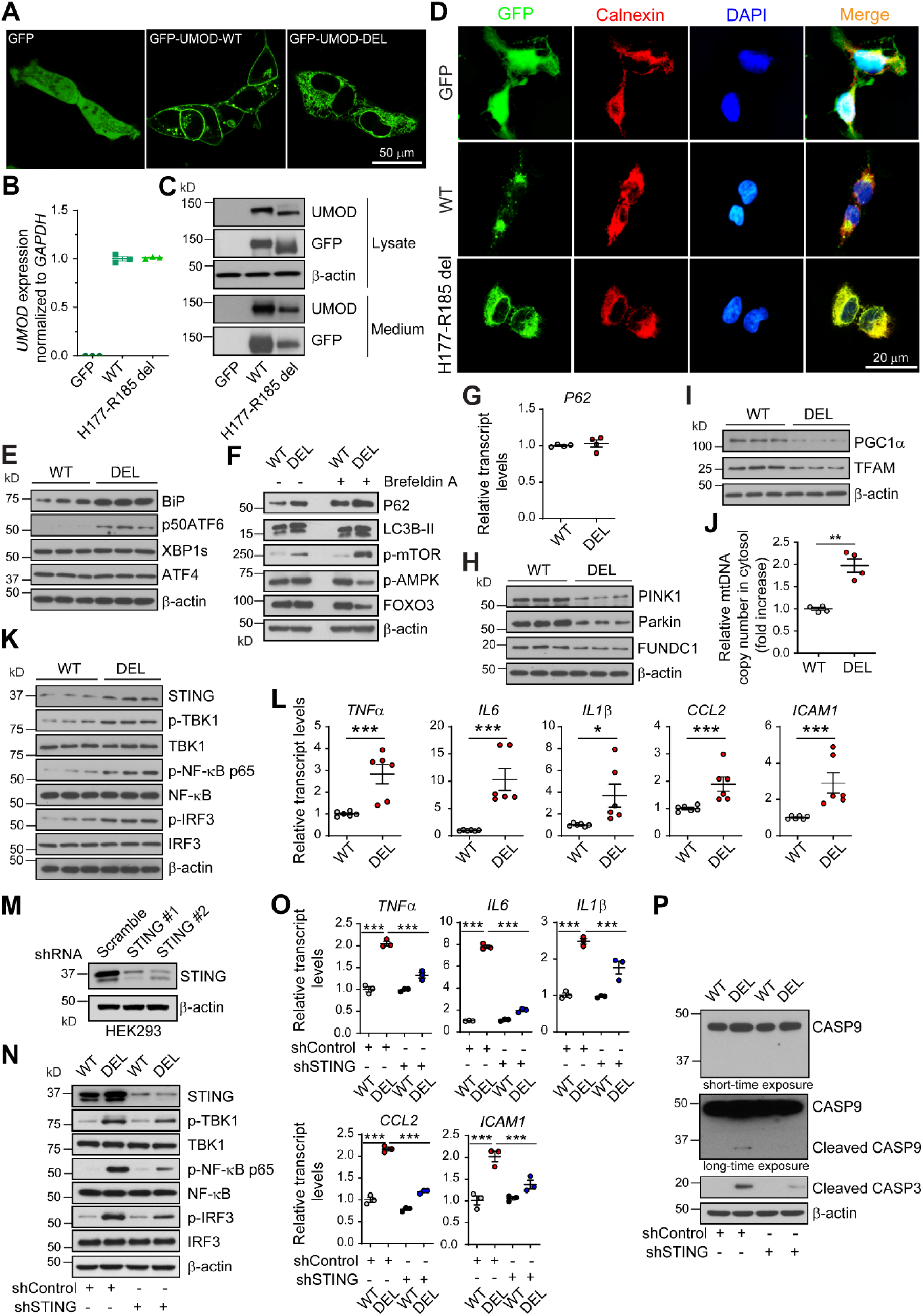
Cytosolic leak of mtDNA activates STING signaling by utilizing stable HEK 293 cell line expressing N-terminal GFP-tagged WT or H177-R185 del UMOD. **(A)** Confocal live imaging showing GFP signal in HEK 293 cells expressing GFP alone, GFP- tagged WT or mutant H177-R185 del uromodulin. Scale bar, 50 μm. **(B)** *UMOD* transcript expression assessed by real-time RT-qPCR. Expression was normalized to GAPDH. The *UMOD* mRNA level in cells expressing GFP alone was shown as negative control. Mean ± SD of fold changes of three independent samples from each clone. **(C)** Cell lysates and media from cells expressing WT or mutant H177-R185 del uromodulin were analyzed by WBs for expression of UMOD and GFP. The WB image shown is representative of three independent experiments. **(D)** IF analysis of GFP (green) and an ER marker, calnexin (red). Nuclei were counterstained with DAPI (blue). Scale bar, 20 μm. **(E)** Cell lysates of WT and DEL cells were analyzed by WBs for protein levels of BiP, p50ATF6, XBP1s and ATF4. N=3 per group. **(F)** Untreated and brefeldin A (10 μg/ml)-treated WT and DEL cells were analyzed by WBs for protein levels of p62, LC3B-II, p-mTOR (S2481), p-AMPK and FOXO3. The cells were exposed to brefeldin A for 4 hours, and then recovered in the cell culture media for 20 hours. The cells were harvested for WB analysis after recovery. The WB image shown is representative of three independent experiments. **(G)**Quantitative PCR analysis for mRNA levels of p62. Gene expression was normalized to GAPDH. Mean ± SD of fold changes of four independent samples from each clone. **(H)**Cell lysates of WT and DEL cells were analyzed by WBs for protein levels of PINK1, Parkin and FUNDC1. N=3 per group. **(I)** Cell lysates of WT and DEL cells were analyzed by WBs for protein levels of PGC1α and TFAM. N=3 per group. **(J)** Cytosolic translocation of mtDNA in WT and DEL cells was quantified by q-PCR. The copy number of mtDNA encoding cytochrome *c* oxidase I was normalized to nuclear DNA encoding 18S ribosomal RNA. Mean ± SD of fold changes. **p<0.01. **(K)**Cell lysates of WT and DEL cells were analyzed by WBs for protein levels of STING, p- TBK1/TBK1, p-NF-κB/NF-κB and p-IRF3/IRF3. N=3 per group. **(L)** Quantitative PCR analysis for mRNA levels of proinflammatory genes downstream of STING/NF-κB signaling. Gene expression was normalized to GAPDH. Mean ± SD of fold changes. *p<0.05; ***p<0.001. **(M)** WB analysis showed knockdown efficacy of STING expression from shSTING1 and shSTING2 vs. a scrambled shRNA control in HEK 293 cells. **(N)** Cell lysates of WT and DEL cells, treated with shControl or shSTING for 48 hours, were analyzed by WBs for protein levels of STING, p-TBK1/TBK1, p-NF-κB/NF-κB and p- IRF3/IRF3. The WB image shown is representative of three independent experiments. **(O)**Quantitative PCR analysis for mRNA levels of proinflammatory genes downstream of STING/NF-κB signaling in cells treated with shControl or shSTING for 48 hours. Gene expression was normalized to GAPDH. Mean ± SD of fold changes. ***p<0.001. **(P)** Cell lysates of WT and DEL cells, treated with shControl or shSTING for 48 hours, were analyzed by WBs for protein levels of cleaved caspase 9 and cleaved caspase 3. The WB image shown is representative of three independent experiments.

DEL cells, as compared to WT cells, exhibited increased expression of BiP and selective activation of the p50ATF6 branch without involvement of XBP1s and ATF4 branches (Figure 5E). DEL cells also showed increased protein levels of P62, LC3B-II and p-mTOR (S2481) (Figure 5F) without transcriptional upregulation of P62 (Figure 5G), as well as decreased expression of PINK1/Parkin/FUNDC1 (Figure 5H) compared to WT cells, indicating insufficient autophagy/mitophagy. The cells were next pulsed for 4 hours with 10 μg/mL brefeldin A (an inhibitor of protein trafficking from the ER to the Golgi apparatus) to increase the accumulation of UMOD in the ER, and then allowed to recover in the growth media for 20 hours. After recovery, the mutant cells evidently displayed increased abundance of P62 and p-mTOR compared to untreated mutant cells (Figure 5F). Moreover, in the brefeldin A-treated cells, expression of p-AMPK and FOXO3 in DEL cells became significantly reduced compared with that in the brefeldin A-treated WT cells (Figure 5F). Meanwhile, decreased protein levels of PGC1α and TFAM, which reflected dysfunctional mitochondrial biogenesis, were noted in DEL cells vs. WT cells (Figure 5I). Together, we conclude that the stable cell model simulates the findings in our ADTKD-*UMOD* mouse model well.

By utilizing the cell model, we isolated cytosolic fractions and directly measured release of mtDNA into the cytosol by normalizing the copy number of mtDNA encoding cytochrome *c* oxidase I to nuclear DNA encoding 18S ribosomal RNA. As shown in Figure 5J, the mtDNA leakage into the cytosol was significantly higher in DEL cells relative to that in the WT cells. Furthermore, consistent with the mouse model, activated STING/p-TBK1/p-NF-κB/p-IRF3 signaling (Figure 5K) and increased transcript levels of inflammatory cytokines, including TNFα, IL6, IL1β, CCL2 and ICAM1 (Figure 5L) were observed in the mutant cells compared with WT cells. We next investigated whether STING knockdown using a lentiviral shRNA-driven approach could dampen cellular inflammation and apoptosis. We tested two shRNAs directed against STING (shSTING). As shown in Figure 5M, expression of STING was robustly reduced in HEK 293 cells by both sequences compared with that by a scrambled shRNA control (shControl). shSTING 1 was chosen for further studies due to the better knockdown efficacy.

Increased STING/p-TBK1/p-NF-κB/p-IRF3 expression (Figure 5N), as well as augmented inflammatory gene expression (Figure 5O) in DEL cells was significantly mitigated by shSTING treatment compared with shControl. As a result, increased caspase-9 and caspase-3 cleavage in DEL cells was attenuated by STING knockdown (Figure 5P). Together, these data directly demonstrate that cGAS-STING signaling is activated by cytosolic leak of mtDNA in H177-R185 del cells, which mediates proinflammatory cytokine production and cell death.

### Loss of MANF in TALs deteriorates autophagy suppression and kidney fibrosis in ADTKD

After we successfully generated the knock-in mouse and cell models resembling ADTKD- *UMOD*, we sought to explore the function of MANF in ER-stressed mutant TALs. Double IF staining of MANF and UMOD revealed increased MANF expression distributed within UMOD^+^ tubules, as early as by 3 weeks of age, in the mutant kidneys (Supplemental Figure 7A). By 16 weeks, MANF upregulation in the TALs expressing the mutant UMOD became conspicuous in both whole-kidney sections by staining (Figure 6A) and protein lysates by WB (Figure 6B). We further confirmed increased MANF level in the UMOD^+^ tubules in the kidney biopsy from a patient carrying *UMOD* H177-R185 del (Figure 6C).

**Figure 6.**
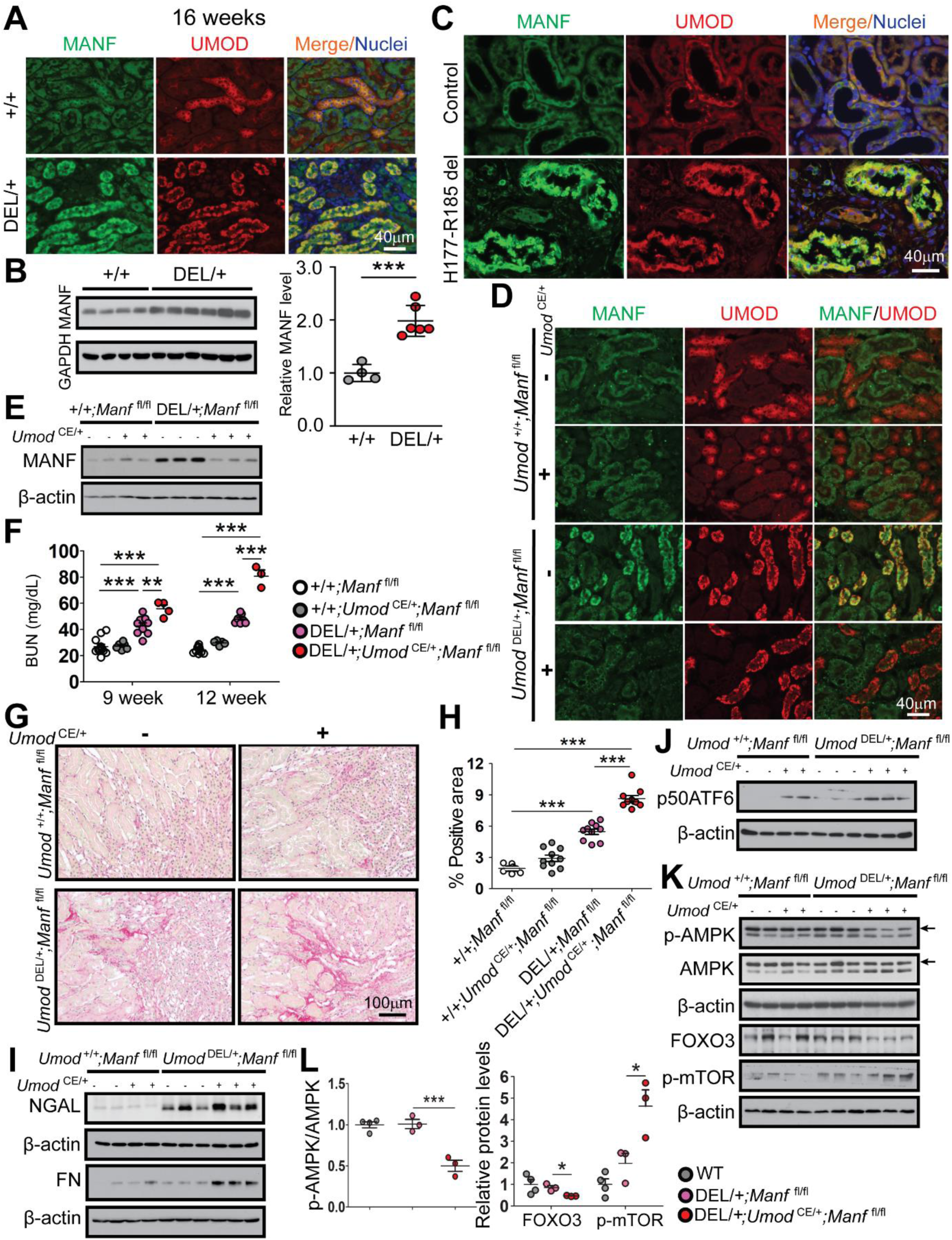
Loss of MANF in TALs deteriorates autophagy suppression and kidney fibrosis in ADTKD-*UMOD*. **(A)** Co-localization of MANF (green) and UMOD (red) on paraffin kidney sections from *Umod ^+/+^* and *Umod* ^DEL/+^ mice at 16 weeks. Nuclei were counterstained with Hoechst 33342 (blue). Scale bar, 40 µm. **(B)** WB analysis of MANF in kidneys harvested from *Umod^+/+^* and *Umod* ^DEL/+^ mice at 16 weeks with densitometry analysis. The average MANF/β-actin ratio in WT mice was set as 1. Mean ± SD (n=4-6 mice/genotype). ***p<0.001. **(C)** Paraffin kidney sections from a patient with p.H177-R185 del and from a normal kidney, stained for MANF (green) and UMOD (red) with nuclei counterstain (blue). Scale bar, 40 µm. **(D)** Paraffin kidney sections from mice of the indicated genotypes were examined by dual IF staining of MANF (green) and UMOD (red) at 9 weeks. Scale bar, 40 µm. **(E)** Representative WB analysis of MANF in kidneys harvested from *Umod* ^+/+^;*Manf* ^fl/fl^, *Umod* ^+/+^;*Umod* ^CE/+^;*Manf* ^fl/fl^, *Umod* ^DEL/+^;*Manf* ^fl/fl^, and *Umod* ^DEL/+^;*Umod* ^CE/+^;*Manf* ^fl/fl^ mice at 12 weeks. **(F)** BUN measurements at 9 and 12 weeks. Mean ± SD (n=3-12 mice/genotype). **p<0.01; ***p<0.001. **(G)**Representative histological images of whole kidney sections stained with Picrosirius red (collagens I/III in red). Scale bar, 100 µm. **(H)**Quantification of Picrosirius Red positive areas. Mean ± SD (n=10 images/genotype). ***p<0.001. **(I)** Representative WBs of NGAL and FN in *Umod^+/+^* and *Umod* ^DEL/+^ kidneys without or with MANF deletion at 12 weeks. **(J)** Representative WB of p50ATF6 in *Umod ^+/+^* and *Umod* ^DEL/+^ kidneys in the presence or absence of MANF in TALs at 12 weeks. **(K)**Representative WBs evaluating p-AMPK (arrow), AMPK (arrow), FOXO3 and p-mTOR (S2481) in *Umod ^+/+^* and *Umod* ^DEL/+^ kidneys without or with MANF deficiency at 12 weeks. **(L)** Densitometry analysis of p-AMPK/AMPK, as well as FOXO3 and p-mTOR normalized to β-actin in kidney lysates of the indicated groups at 12 weeks. The average p-AMPK/AMPK, FOXO3 or p-mTOR/β-actin ratio in WT mice was set as 1. Mean ± SD (n=3-4 mice/group). *P<0.05; ***P<0.001. **(D-L)** All mice were given TAM at 5 weeks of age.

Next, we studied the biological function of MANF in ADTKD-*UMOD*. To specifically ablate MANF from UMOD^+^ TALs in *Manf* ^fl/fl^ mice, we used a tamoxifen (TAM)-dependent *Umod*- driven Cre recombinase line (*Umod* ^IRES CRE-ERT^^2^, hereafter referred to as *Umod* ^CE^), in which IRES-CRE-ERT2 was knocked into the 3’ untranslated region (UTR) near the stop codon of the *Umod*. Their offspring, *Umod* ^DEL/+^;*Manf* ^fl/fl^ and *Umod* ^DEL/+^;*Umod* ^CE/+^;*Manf* ^fl/fl^, as well as their WT control littermates *Umod* ^+/+^;*Manf* ^fl/fl^ and *Umod* ^+/+^;*Umod* ^CE/+^;*Manf* ^fl/fl^, was given TAM for three doses (3 mg/mouse), every other day at 5 weeks of age. Subsequently, the TAM-administered cohorts of mice were followed until 12 weeks. The specific removal of MANF from TAL cells after TAM treatment was confirmed by PCR of genomic DNA isolated from TALs (Supplemental Figure 7B). As expected ^18^, the amplified band of 1300 bp contained exon 3 of *Manf* and Frt- and LoxP-sites in *Umod* ^+/+^;*Manf* ^fl/fl^ and *Umod* ^DEL/+^;*Manf* ^fl/fl^ mice, while 531 bp represented the targeted *Manf* ^-/-^ lacking the exon 3 in *Umod* ^+/+^;*Umod* ^CE/+^;*Manf* ^fl/fl^ and *Umod* ^DEL/+^;*Umod* ^CE/+^;*Manf* ^fl/fl^ littermates (Supplemental Figure 7B). In line with the PCR results, this manipulation led to a significant reduction of MANF protein levels in the TALs producing mutant UMOD, as evidenced by co-IF staining of MANF and UMOD (Figure 6D) and immunoblot analysis of MANF (Figure 6E). At 1 month after administration of TAM, we first observed a significant elevation of BUN in *Umod* ^DEL/+^;*Umod* ^CE/+^;*Manf* ^fl/fl^ mice compared with *Umod* ^DEL/+^;*Manf* ^fl/fl^ mice at 9 weeks (Supplemental Figure 7C and Figure 6F). At 12 weeks, the accelerated progression of kidney disease in the mutant mice deficient of MANF in the TALs became much more pronounced (Figure 6F). Consistent with exacerbation of the kidney function after depletion of MANF in TALs, Picrosirius red staining revealed marked increases in collagens I and III in the corticomedullary junction area (Figure 6, G-H), and WB showed an increase in NGAL and FN in *Umod* ^DEL/+^;*Umod* ^CE/+^;*Manf* ^fl/fl^ kidneys compared with *Umod* ^DEL/+^;*Manf* ^fl/fl^ kidneys at 12 weeks (Figure 6I).

Mechanistically, MANF deficiency in TALs was linked to an increased expression of p50 ATF6, on both WT and *Umod* ^DEL/+^ background (Figure 6J). We also observed that MANF deletion in TALs resulted in a further failure in autophagy, as demonstrated by decreased levels of the positive autophagy regulators p-AMPK (arrow) and FOXO3, as well as increased level of the negative regulator p-mTOR (S2481) in the whole-kidney lysates from *Umod* ^DEL/+^;*Umod* ^CE/+^;*Manf* ^fl/fl^ mice vs. *Umod* ^DEL/+^;*Manf* ^fl/fl^ mice at 12 weeks (Figure 6, K and L). Together, these data support that MANF upregulation in ER-stressed TAL cells is indispensable for maintaining autophagic activity, and that loss of MANF in mutant TALs exacerbated autophagy inhibition, further intensifying ER stress and kidney injury in ADTKD-*UMOD*.

### Transgenic (Tg) tubular MANF expression stimulates autophagy and clearance of mutant UMOD in the mouse model of ADTKD-*UMOD*

We next asked whether MANF overexpression can enhance autophagy and play a therapeutic role in ADTKD-*UMOD*. To this end, we generated inducible MANF Tg (TET-MANF) mice, in which the full-length cDNA of human MANF is controlled by a (TetO)7/CMV promoter containing 7 tetracycline-response elements (TREs) (Supplemental Figure 8Ab). Next, TET- MANF mice were crossed with Pax8-reverse tetracycline-controlled transcriptional activator (rtTA) mice (Pax8-rtTA mice, abbreviated to P8TA hereafter), in which the pan-tubular specific promoter Pax8 directs expression of rtTA in all renal tubular epithelial cells (Supplemental Figure 8Aa). In the presence of tetracycline derivative, doxycycline (DOX), rtTA binds to TRE and initiates transcription of the MANF cDNA in renal epithelial cells (Supplemental Figure 8A). When TET-MANF was bred to homozygous P8TA/P8TA mice, two genotypes were generated and abbreviated to MANF/P8TA and P8TA. The offspring was induced with DOX at 4 weeks. As shown in Figure 7A, by 2 weeks after the DOX induction, urinary MANF excretion was easily detected in MANF/P8TA, but not in P8TA mice. Q-PCR of MANF using a pair of primers shared in both mouse and human MANF genomic sequences showed a 2.33 ± 0.3 fold of increase of MANF in the DOX-treated bi-transgenic compared with the single Tg mice after 4 weeks administration of DOX (Figure 7B). At the protein level, the double Tg mice exhibited increased abundance of MANF compared to single Tg mice with 4 weeks of DOX treatment (Figure 7C). Moreover, double IF staining clearly demonstrated that MANF was induced in both UMOD^+^ tubules (Figure 7D, red arrows) and UMOD^-^ tubules (Figure 7D, white arrows) in a perinuclear distribution pattern. Co-IF staining of MANF with the proximal tubular marker, lotus tetragonolobus lectin (LTL) confirmed that proximal tubules also overexpress MANF with DOX treatment in MANF/P8TA mice (Supplemental Figure 8B).

**Figure 7.**
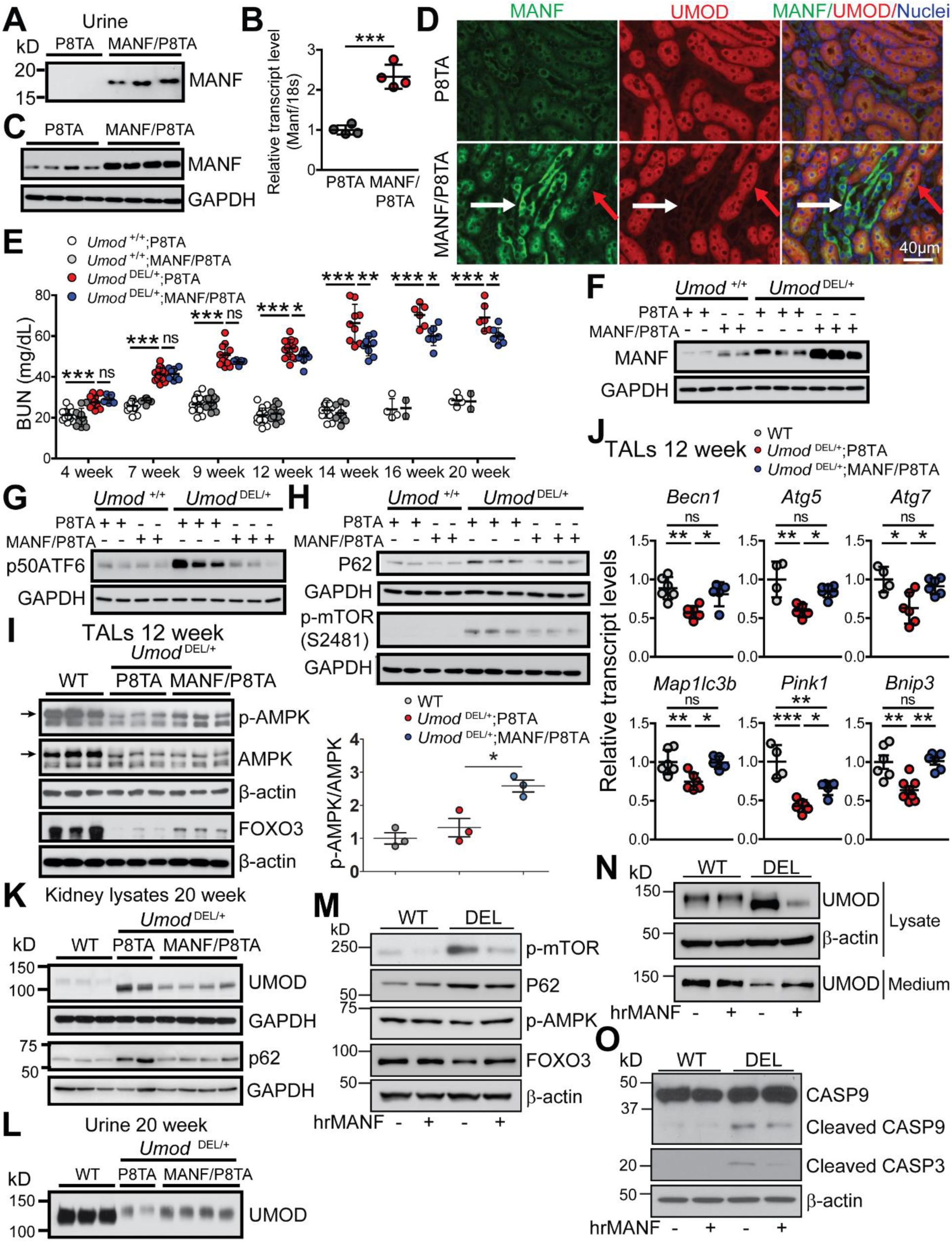
MANF treatment *in vivo* and *in vitro* stimulates autophagy in ADTKD. **(A)** Representative WB analysis of MANF in urine specimens collected from P8TA and MANF/P8TA mice treated with DOX for two weeks. The urinary MANF excretion was normalized to 4 μg of urine creatinine excretion. n = 3 mice/genotype. **(B)** Quantitative RT-PCR analysis of relative transcript level of MANF in DOX-treated P8TA and MANF/P8TA kidneys for 4 weeks. Gene expression was normalized to 18s. Mean ± SD (n=4 mice/genotype). ***p<0.001. **(C)** Kidneys from P8TA and MANF/P8TA mice fed with DOX starting from 4 weeks of age were harvested after 4 wks of DOX administration. WB analysis of MANF expression in kidney lysates from the indicated groups. n=4 mice/genotype. **(D)** IF staining of MANF (green), UMOD (red) and Hoechst 33342 (blue) on kidney paraffin sections of the indicated groups after 4 weeks of DOX administration. Red arrows point to UMOD^+^ tubules and white arrows point to UMOD^-^ tubules. Scale bar, 40 µm. **(E)** BUN measurements in the indicated groups over a 20-week period. Mean ± SD (n=5-19 mice/genotype). ns, not significant; *p<0.05; **p<0.01; ***p<0.001. **(F)** Representative WB from *Umod ^+/+^* and *Umod* ^DEL/+^ kidneys in the presence or absence of tubular MANF overexpression for levels of MANF at 14 weeks. **(G)**Representative WB from *Umod ^+/+^* and *Umod* ^DEL/+^ kidneys in the presence or absence of tubular MANF overexpression for levels of p50ATF6 at 14 weeks. **(H)**Representative WBs from the kidneys of the indicated groups for levels of p62 and p- mTOR (S2481) at 14 weeks. **(I)** Representative WBs from *Umod ^+/+^* and *Umod* ^DEL/+^ TAL cells with or without tubular MANF overexpression for levels of p-AMPK (arrow), AMPK (arrow) and FOXO3 at 12 weeks with densitometry analysis of p-AMPK/AMPK. The average p-AMPK/AMPK ratio in WT mice was set as 1. Mean ± SD (n=3 mice/group). **(J)** Quantitative PCR of isolated primary TALs from the indicated groups at 12 weeks for autophagy and mitophagy-related genes. Mean ± SD (n=4-8 mice/genotype). ns, not significant; *p<0.05; **p<0.01; ***p<0.001. **(K)**Representative WBs from kidneys of the indicated groups for levels of UMOD and P62 at 20 weeks. **(L)** Representative WB from urine of the indicated groups for levels of WT UMOD at 20 weeks. The urinary UMOD excretion was normalized to 4 μg of urine creatinine excretion. **(M-O)** Stably transduced HEK 293 cells expressing WT or H177-R185 del UMOD were pulsed with 10 μg/mL brefeldin A for 4 hours, and then allowed to recover in the growth media for 20 hours. After recovery, WT and DEL cells were treated with human recombinant MANF (hrMANF, 5 μg/ml) for 24 hours. Cell lysates of WT and DEL cells treated without or with hrMANF were analyzed by WBs for protein levels of p-mTOR, P62, p-AMPK and FOXO3 **(M)**, UMOD **(N)**, and cleaved caspase 9 and caspase 3 **(O)**. The media of WT and DEL cells treated without or with hrMANF were also analyzed by WB for UMOD secretion **(N)**. The WB image shown is representative of three independent experiments. **(A-L)** All mice were given DOX.

After we had characterized a successful tubular induction of MANF with DOX administration, TET-MANF mice were crossed with *Umod* ^DEL/+^;P8TA/P8TA mice. The 4 week- old offspring was started with DOX after the onset of disease. BUN was monitored longitudinally in the offspring until 20 weeks. We observed a statistically significant BUN decrease in *Umod* ^DEL/+^;MANF/P8TA vs. *Umod* ^DEL/+^;P8TA mice from 12 to 20 weeks (Figure 7E). We further investigated the molecular mechanisms underlying the protective effect of MANF by utilizing the kidney lysates at 14 weeks. First, we confirmed that kidneys from DOX- treated MANF/P8TA mice exhibited significantly increased MANF levels compared to kidneys from DOX-treated P8TA mice, either on the *Umod* ^+/+^ or *Umod* ^DEL/+^ background (Figure 7F).

ATF6 activation in *Umod* ^DEL/+^;P8TA kidneys was substantially suppressed by the tubular upregulation of MANF (Figure 7G). Next, we looked at the effect of MANF on autophagic activity. With enhanced expression of MANF in the *Umod* ^DEL/+^;MANF/P8TA kidneys, the P62 level was decreased compared with that in the *Umod* ^DEL/+^;P8TA kidneys, indicating increased autophagic activity (Figure 7H). In addition, MANF upregulation in *Umod* ^DEL/+^;MANF/P8TA kidneys attenuated expression of p-mTOR (S2481) in *Umod* ^DEL/+^;P8TA kidneys, thus promoting autophagic flux (Figure 7H). Moreover, increased levels of p-AMPK (arrow) and FOXO3 by MANF overexpression in isolated primary mutant TALs at 12 weeks corroborated the findings in the whole-kidney lysates at 14 weeks (Figure 7I). Transcript analysis of TALs from the indicated genotypes further supports that decreased mRNA levels of key autophagy/mitophagy-related genes, including *Becn1*, *Atg5*, *Atg7*, *Map1lc3b* and *Bnip3*, in *Umod* ^DEL/+^;P8TA TAL cells were restored by MANF upregulation toward WT levels (Figure 7J)*. Pink1* exhibited partial restoration by MANF overexpression (Figure 7J). At 20 weeks of age, a marked decrease of mutant UMOD and P62 in the DOX-treated *Umod* ^DEL/+^;MANF/P8TA kidneys was observed compared with those in the DOX-treated *Umod* ^DEL/+^;P8TA kidneys (Figure 7K). Meanwhile, urinary excretion of WT UMOD was considerably increased in the DOX-treated *Umod* ^DEL/+^;MANF/P8TA mice compared with DOX-treated *Umod* ^DEL/+^;P8TA mice (Figure 7L).

To directly demonstrate that the secreted MANF acts as an autophagy activator, we treated the WT and mutant cells producing H177-R185 del UMOD with human recombinant MANF (hrMANF, R&D). The cells were shortly exposed to low dose brefeldin A (10 μg/mL) for 4 hours to promote UMOD retention in the ER. After recovery in the culture media for 20 hours, the cells were treated with 5 μg/mL of hrMANF for 24 hours. In contrast to DEL cells in the absence of MANF, hrMANF-treated DEL cells manifested a decrease in the abundance of p-mTOR (S2481) and P62, and an increase in p-AMPK and FOXO3 (Figure 7M). Due to the increased autophagy flux, hrMANF treatment enhanced autophagic degradation of mutant UMOD in the DEL cell lysates and improved secretion of WT UMOD into the cell culture medium in DEL cells (Figure 7N), consistent with the *in vivo* findings (Figure 7, K and L). Consequently, treatment with hrMANF abolished activation of caspase-9 and caspase-3 in the mutant cells (Figure 7O). Collectively, these studies suggest that secreted MANF treatment promotes autophagic activity and removal of misfolded UMOD in ADTKD-*UMOD*.

### MANF overexpression in renal tubules restores mitochondrial homeostasis, suppresses STING activation, and ameliorates kidney injury and fibrosis in ADTKD-*UMOD*

Given that MANF upregulation stimulates expression of p-AMPK, the guardian of mitochondrial homeostasis and metabolism, we therefore reasoned that tubular MANF overexpression may improve mitochondrial function in ADTKD-*UMOD* mouse model. Prompted by increased transcriptional expression of *Pink1* and *Bnip3* by MANF (Figure 7J), we first investigated the effect of MANF on mitophagy. Increased protein abundance of PINK1/Parkin/BNIP3L (arrow) by MANF overexpression in isolated primary mutant TAL cells at 12 weeks concurred with the notion that MANF promoted mitophagy (Figure 8A). In addition, at 12 weeks, WB revealed increased expression of PGC1α in the TALs purified from DOX-treated *Umod* ^DEL/+^;MANF/P8TA kidneys compared with TALs from DOX-treated *Umod* ^DEL/+^;P8TA kidneys (Figure 8B). In alignment with increased PGC1α expression, mutant TALs with MANF overexpression exhibited increased levels of mitochondrial gene transcripts, including *mt-Nd4, mt-Co1, mt-CytC and mt-Rnr2* (Figure 8C), and ETC proteins (Figure 8D) relative to mutant TALs without MANF transgenic expression at 16 weeks. Finally, the high resolution respirometry showed that mitochondrial respiration was significantly higher in the MANF- overexpressing mutant TAls as compared to mutant TALs at 16 weeks (Figure 8E).

**Figure 8.**
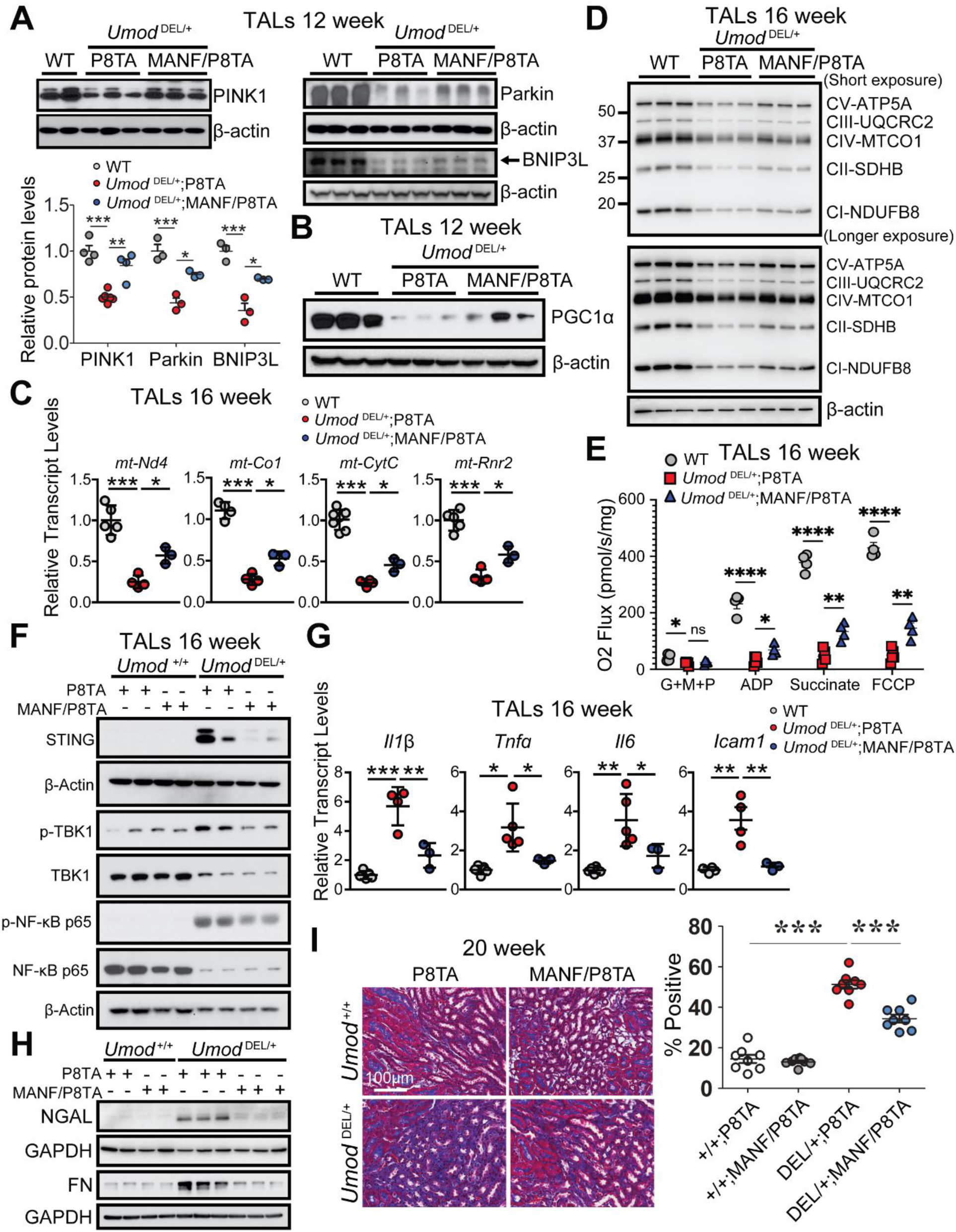
Renal tubular upregulation of MANF promotes mitophagy and improves mitochondrial biogenesis, leading to abrogation of STING activation and kidney fibrosis in ADTKD-*UMOD*. **(A)** Representative WBs of PINK1, Parkin and BNIP3L (arrow) in TALs isolate from the kidneys of the indicated groups at 12 weeks with densitometry analysis. The average PINK1, Parkin, or BNIP3L//β-actin ratio in WT mice was set as 1. Mean ± SD (n=3-6 mice/group). *p<0.05; **p<0.01; ***p<0.001. **(B)** Representative WB of PGC1α in TALs from the kidneys of the indicated groups at 12 weeks. n=3 mice/group. **(C)** Transcript analysis of *mt-Nd4, mt-Co1, mt-CytC and mt-Rnr2* from TALs of the indicated genotypes at 16 weeks. Mean ± SD (n=3-6 mice/genotype). *p<0.05; ***p<0.001. **(D)** Representative WBs of ETC expression in TALs of the indicated genotypes at 16 weeks. n=3 mice per genotype. **(E)** Assessment of mitochondrial function using an OROBOROS Oxygraph system in permeabilized TALs of the indicated genotypes at 16 weeks following sequential additions of glutamate, malate and pyruvate (G+M+P); adenosine diphosphate (ADP); succinate and FCCP. Mean ± SD (n=4-5 mice/genotype). *p<0.05, **p<0.01, ****p<0.0001. **(F)** Representative WBs of STING, p-TBK1/TBK1 and p-NF-κB/NF-κB in TALs isolate from the kidneys of the indicated genotypes at 16 weeks. **(G)**Transcript analysis of *Il1b, Tnfα*, *Il6* and *Icam1* from TALs of the indicated genotypes at 16 weeks. Mean ± SD (n=3-5 mice/genotype). *p<0.05; **p<0.01; ***p<0.001. **(H)**Representative WBs of NGAL and FN in the kidneys of the indicated genotypes at 20 weeks. **(I)** Representative Gomori’s Trichrome staining of paraffin kidney sections from the kidneys of the indicated groups at 20 weeks with quantification. Scale bar, 100 µm. Mean ± SD (n=8 images/genotype). ***p<0.001. **(A-I)** All mice were given DOX.

Correlating with the effect on improving mitochondrial health, tubular upregulation of MANF ameliorated STING activation in *Umod* ^DEL/+^;P8TA TALs, as evidenced by decreased protein levels of STING, p-TBK1 and p-NF-κB in *Umod* ^DEL/+^;MANF/P8TA TALs at 16 weeks (Figure 8F). We further conducted q-PCR analysis of downstream targets of STING/NF-κB signaling on isolated TAL cells at 16 weeks. The mRNA levels confirmed that upregulation of *Il1b*, *Tnfa*, *Il6* and *Icam1* gene expression in mutant UMOD-expressing tubular epithelium was blunted by MANF overexpression (Figure 8G). Consequently, enhanced expression of MANF in renal tubules repressed expression of the tubular injury marker NGAL, and fibrosis marker FN at 20 weeks (Figure 8H). In line with immunoblot results, Trichrome staining further confirmed that MANF overexpression in renal tubules significantly blocked fibrosis in the mutant kidneys (Figure 8I). In summary, our data have convincingly demonstrated that tubular targeted MANF upregulation promotes mitophagy, improves mitochondrial biogenesis and OXPHOS, and mitigates STING activation, leading to attenuation of tubular injury and fibrosis in ADTKD- *UMOD*.

### MANF upregulation in renal tubules slows down CKD progression in *Umod* ^C147W/+^ mice

To explore whether MANF is also a biotherapeutic protein for ADTKD caused by other *UMOD* mutations, we utilized *Umod* ^C147W/+^ mice ^9^, which harbor a human *UMOD* missense mutation and develop CKD and renal fibrosis more slowly than our deletion mutation mice. TET-MANF were bred with *Umod* ^C147W/+^;P8TA/P8TA mice and their offspring, including *Umod*^C147W/+^;P8TA, *Umod* ^C147W/+^;MANF/P8TA together with their WT littermates, *Umod* ^+/+^;P8TA, *Umod* ^+/+^;MANF/P8TA were induced with DOX at 6 weeks, when BUN did not exhibit a significant elevation between the mutants and WT littermates. Serial BUNs were monitored over the 24-week course. As shown in Supplemental Figure 9A, BUN became significantly lower in the DOX-treated *Umod* ^C147W/+^;MANF/P8TA mice compared to DOX-treated *Umod* ^C147W/+^;P8TA mice since 10 weeks of age. Light microscopic examination of Picrosirius red-stained kidney sections revealed less renal fibrosis at 24 weeks (Supplemental Figure 9B) in the *Umod* ^C147W/+^;MANF/P8TA kidneys vs. *Umod* ^C147W/+^;P8TA kidneys at 24 weeks. Together, the data obtained in another line of *Umod* mutant mice support that the renal protective effect of MANF may not be restricted to an individual *Umod* mutation. Further mechanistic studies will be performed to determine whether MANF also promotes autophagy/mitophagy and protects mitochondrial function in *Umod* ^C147W/+^ mice.

## DISCUSSION

In summary, by generating a mouse model that resembles ADTKD-*UMOD* carrying a prevalent human mutation, and by employing both loss- and gain-of-function studies of MANF, our work has discovered that MANF is an important regulator of autophagy/mitophagy and mitochondrial homeostasis of mutant TALs through activation of p-AMPK in ADTKD-*UMOD*, thereby mitigating cGAS/STING activation and promoting autophagic degradation of mutant UMOD (Figure 9). For the first time, our study uncovers novel mechanisms of MANF action and provides a new foundation for pharmacological strategies that target MANF, which may have profound therapeutic potential to ameliorate kidney fibrosis and slow down kidney function decline in ADTKD-*UMOD*.

**Figure 9.**
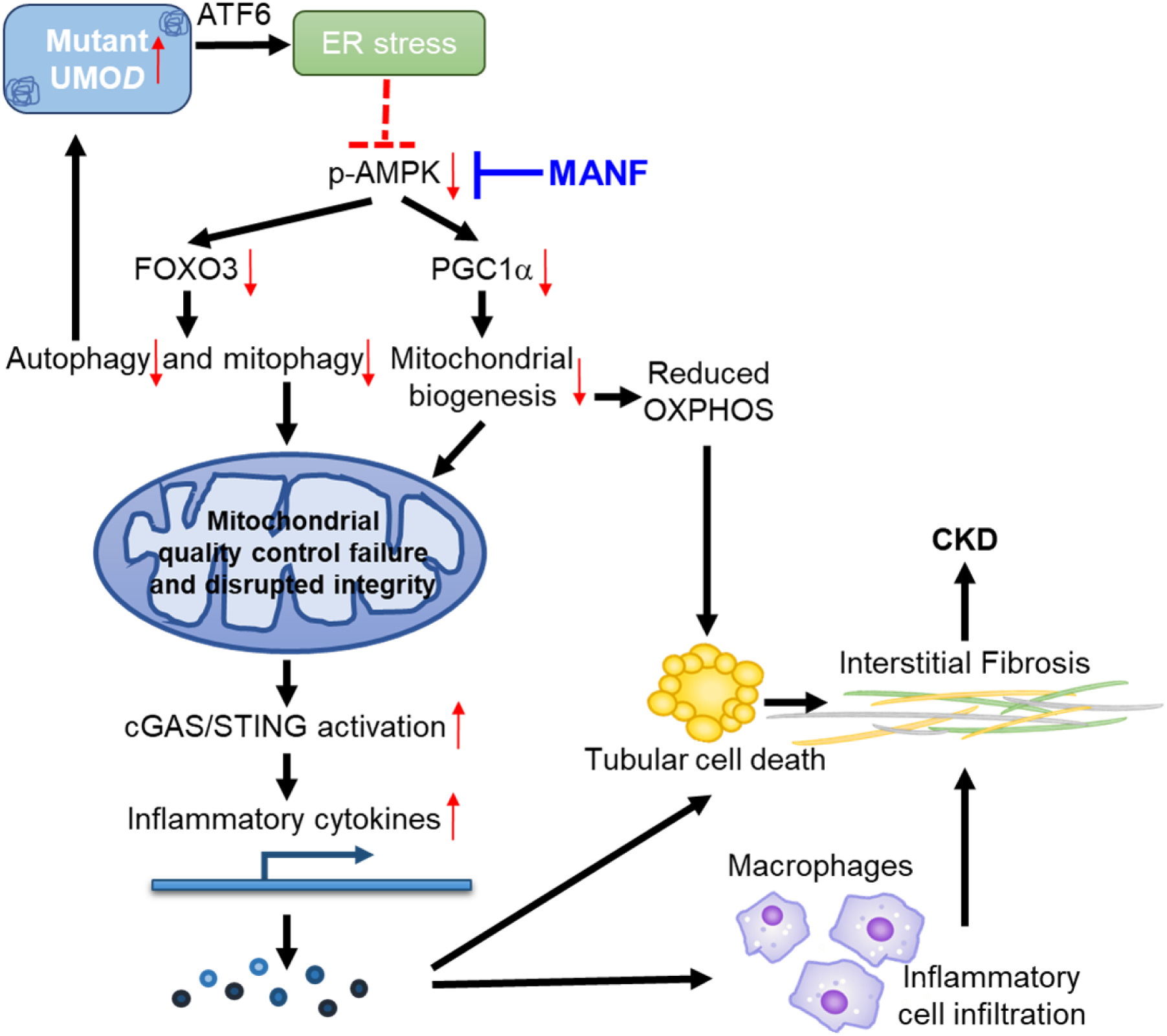
Modulation of kidney fibrosis in ADTKD-*UMOD* by MANF. Mutant UMOD triggers ER stress response, which may directly or indirectly inhibit AMPK activation. Tubular overexpression of MANF after the onset of ADTKD promotes autophagy/mitophagy via activation of p-AMPK-FOXO3 signaling, and enhances mitochondrial biogenesis via p-AMPK-PGC1α axis. Moreover, elimination of dysfunctional mitochondria and restoration of mitochondrial homeostasis through MANF upregulation abrogate STING activation and mitigate tubular pathological inflammation. Finally, increased autophagic clearance of mutant UMOD by MANF induction would further stop the malicious cycle between impaired autophagy and ER stress intensification, eventually protecting kidney function in ADTKD-*UMOD*.

ADTKD is a genetically heterogeneous disease, caused by mutations in *UMOD*, *MUC1* (mucin 1), *HNF1B* (hepatocyte nuclear factor 1β), *REN* (renin) and *SEC61A1* (α1 subunit of translocon 61). Of all 135 *UMOD* mutations identified so far, with the exception of 6 in-frame deletions, most are missense mutations and ∼ 50% of known *UMOD* mutations affect cysteine leading to protein misfolding ^35^. In the current study, we generated the first *Umod* in-frame deletion mouse carrying the most common deletion mutation in human patients. Although cysteine is not present in the deleted 9 amino acids, YETLTEYWR (p.Tyr178_Arg186del), the UMOD deletion mutation activates ER stress, as other missense mutations, presumably due to protein misfolding and aggregation. Notably, different UPR branches were involved in the different *Umod* mutation mouse models. For example, ATF4 and XBP1s are activated (ATF6 was not checked) in *Umod* ^C147W/+^ mice ^9^, whereas these two branches are not upregulated in our deletion mutation mice. It is also interesting to note that in a recently generated ADTKD-*MUC1* mouse model that recapitulates a frame shift mutation in *Muc1* (*Muc1*-fs mice), ATF6 is the prominent UPR branch that regulates ER stress response in response to the MUC-1 protein aggregates while with minimal transcriptional changes in the other two branches ^36^. The function of ATF6 can be adaptive or maladaptive, depending on the different contexts, and few studies have been published regarding the role of ATF6 in the kidney disease. It has been shown that overexpression of p50ATF6 in a proximal tubular cell line decreases the expression of peroxisome proliferator-activated receptor-α (PPAR-α), the master regulator of lipid metabolism, to repress β-oxidation of fatty acid, and thus to cause excessive lipid droplet formation. In contrast, ATF6 ^-/-^ mice have sustained expression of PPAR-α and less tubular lipid accumulation following ischemia-reperfusion injury, leading to less renal fibrosis ^37^. That augmented kidney expression of p50ATF6 through MANF deletion and attenuated expression of ATF6 via MANF overexpression are associated with the opposite kidney functional outcomes in our mouse model might suggest that the effect of ATF6 activation is deleterious in chronic ER-stressed TAL cells. Whether ATF6 directly regulates AMPK activity and thus autophagy/mitophagy, and whether reprogramming the ER proteostasis environment through ATF6 manipulation can impact the disease outcome in ADTKD-*UMOD* are important questions that remain to be answered.

A functional mitophagy system acts as a scavenger of damaged mitochondria and thereby maintains mitochondrial homeostasis. To enable formation of mitophagosomes in mitophagy, mitochondrial priming is regulated by both ubiquitin-dependent PINK1/Parkin pathway and ubiquitin-independent mitophagy receptors, including BNIP3/BNIP3L/FUNDC1 ^25^. It has been reported that PINK1, Parkin or BNIP3 deficiency exacerbates ischemic/reperfusion-, cisplatin-, contrast medium-, or sepsis-induced AKI ^38–42^, as well as unilateral ureter obstruction-induced CKD ^43, 44^. Concurring with the notion that mitophagy plays a protective role in proximal tubular cells under pathogenic conditions, our study is the first to further illuminate the important therapeutic implication of MANF as a mitophagy/autophagy activator in ADTKD-*UMOD*, which can also directly promote autophagic clearance of mutant UMOD. Mechanistically, MANF exerts its mitophagy/autophagy-promoting effect by enhancing p-AMPK-FOXO3 signaling. Recently, we have identified that MANF binds to and antagonizes cell surface receptor neuroplastin to modulate its anti-inflammatory effect ^45^. More studies will be required to understand whether neuroplastin is also involved in regulating AMPK and mitophagy/autophagy in ADTKD.

To our knowledge, this study also provides the first evidence that disrupted mitochondrial integrity due to impaired mitophagy and mitochondrial biogenesis in mutant TALs contributes to STING activation and renal fibrosis in ADTKD-*UMOD*. In alignment with recent studies that mitochondrial damage causes STING activation, renal inflammation and tubular cell death in CKD ^32^ and acute kidney injury (AKI) ^33^, our study highlights the importance of STING signaling in ADTKD-*UMOD*. Furthermore, tubular MANF overexpression directly suppresses STING activation in the mutant TALs.

To date, there is no disease-specific therapy to treat or halt the disease progression of ADTKD until the patients reach end-stage renal disease. It has been shown that blocking TNFα- mediated inflammation by using a soluble TNF receptor inhibitor slows disease progression in *Umod* ^C147W/+^ mice ^9^. Given the hierarchical order from mitochondrial quality control failure for the eventual activation of proinflammatory cytokines in our deletion mutation mice, we envision that treatment with the upstream p-AMPK enhancer MANF, which can restore homeostasis of dysfunctional mitochondria, might be more effective in ADTKD-*UMOD*. MANF is an 18kD protein and recombinant MANF is readily available. In the future, we will continue to test the therapeutic application of MANF as a novel strategy for the treatment of ADTKD patients caused by various gene mutations. In addition, whether MANF can treat other proteinopathies resulting from mutant protein aggregates and altered proteostasis, such as Alzheimer’s disease ^46^, amyotrophic lateral sclerosis ^2^ and retinitis pigmentosa ^47^, is of great interest for the future investigation.

## METHODS

### Antibodies and primers (Tables 1-4)

#### shRNAs and reagents

Lentiviral vector pLKO.1 was used for generating shRNAs targeting STING. Target sequences were selected as follows: shSTING#1, 5′-GCATTACAACAACCTGCTACG-3′; shSTING#2, 5′- GCATCAAGGATCGGGTTTACA-3′.

**Table 1.**
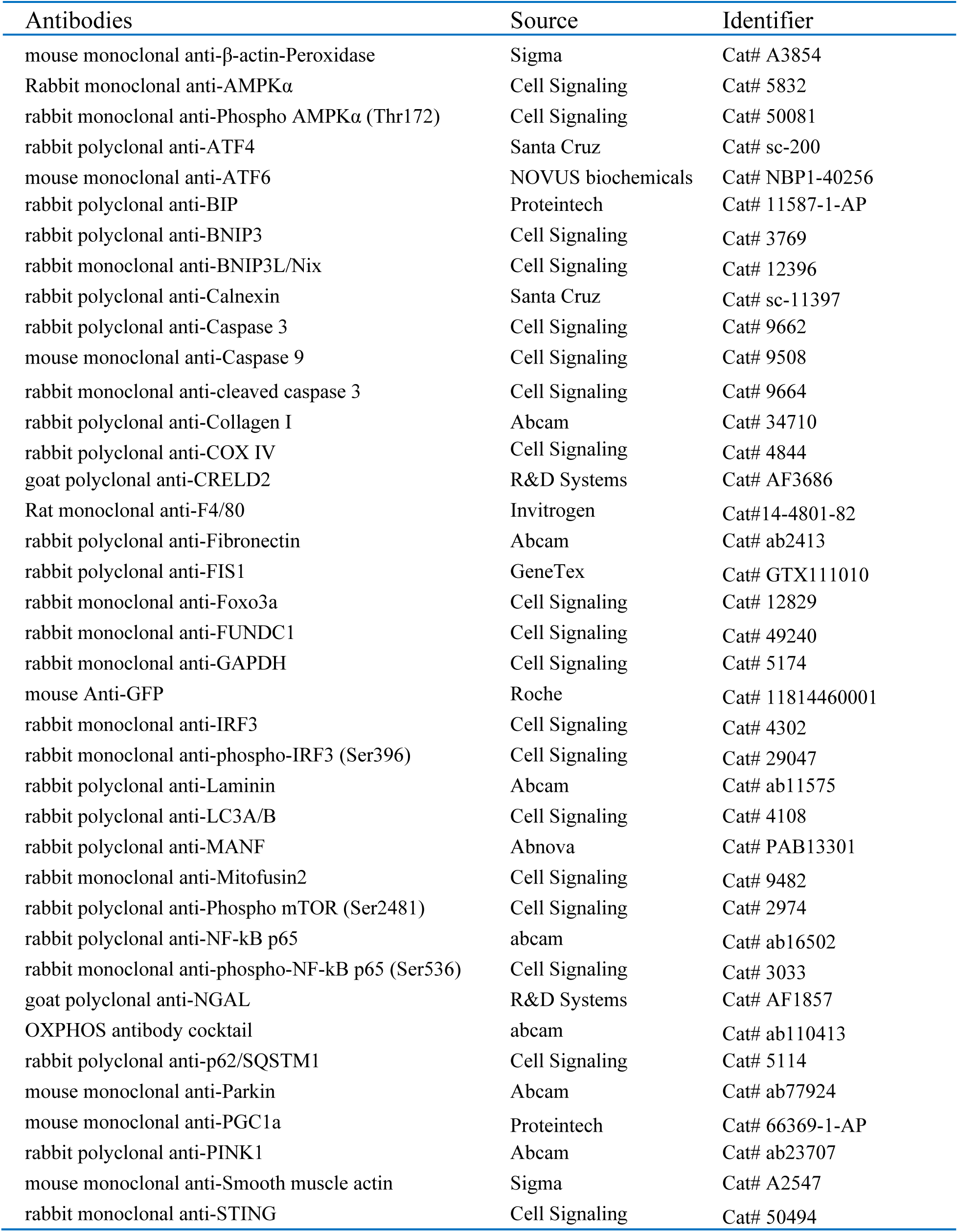
List of Primary Antibodies.

**Table 2.**
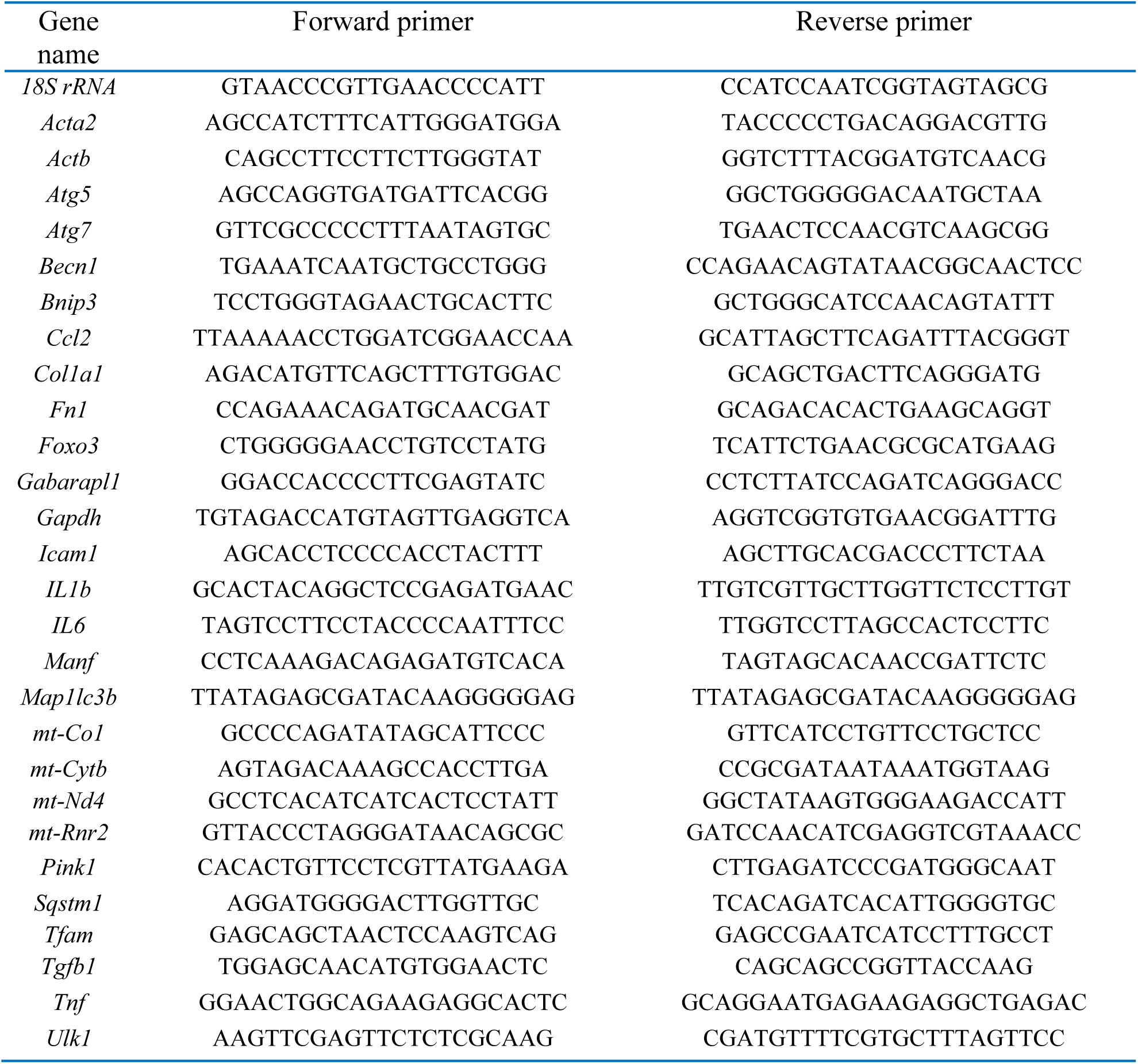
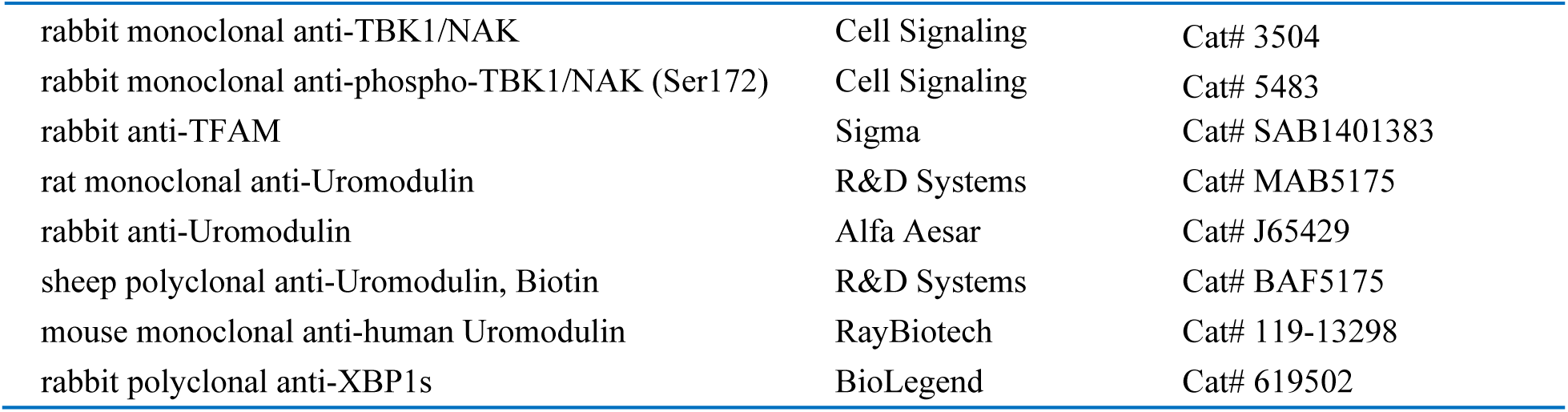
Q-PCR primers for kidneys and TALs.

**Table 3.**
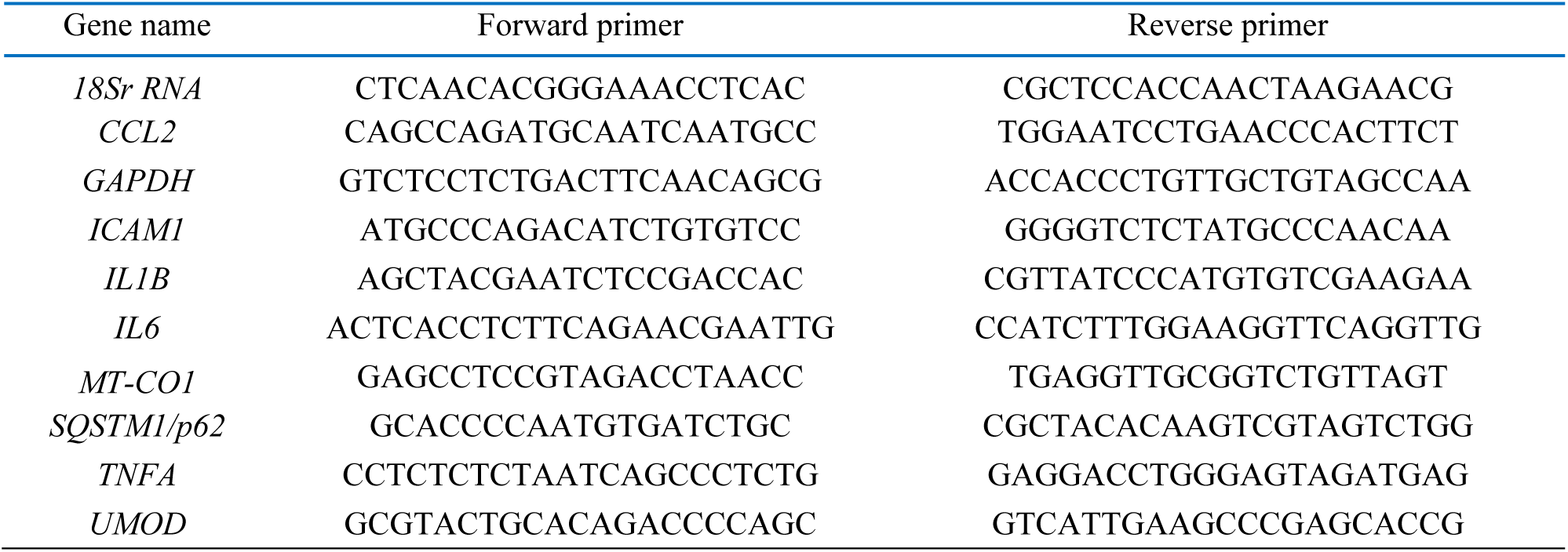
Q-PCR primers for stably transduced HEK 293 cells.

**Table 4.**
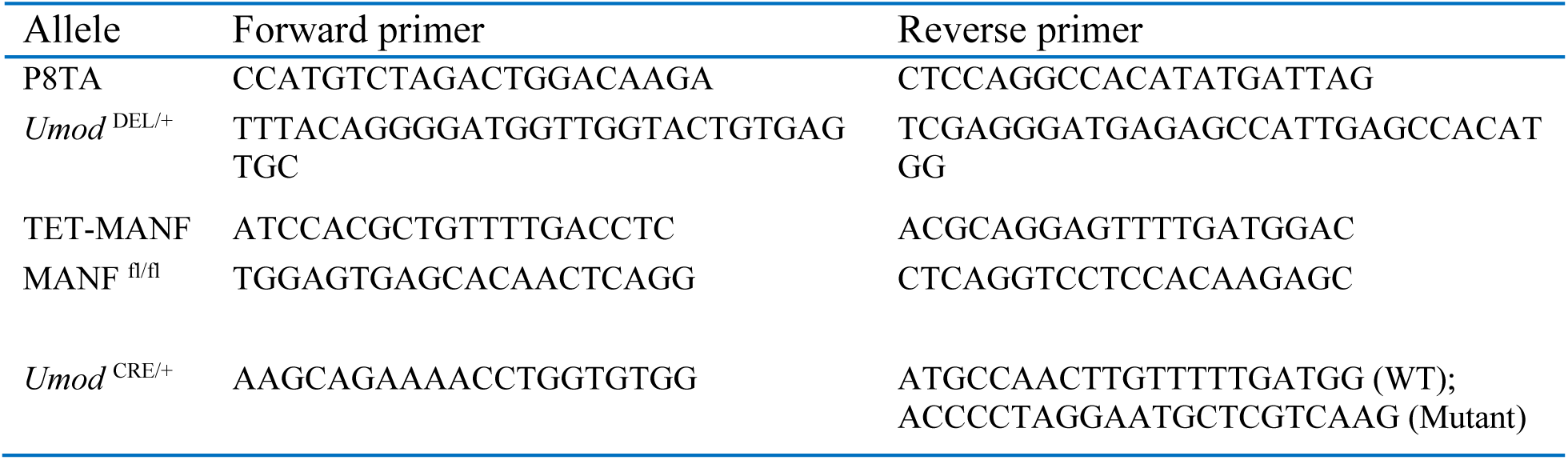
PCR primers for genotyping.

Brefeldin A was purchased from Sigma (St louis, MO, CAT B6542), human recombinant MANF protein was from R&D (Minneapolis, MN, CAT 3748-MN-050), TAM was from Sigma (CAT T5648) and doxycyline food (1500 ppm, irradiated) was from El Mel (St. Charles, MO, CAT 1814935)

#### Animals

Animal experiments conformed to the National Institutes of Health Guide for the Care and Use of Laboratory Animals were approved by the Washington University Animal Studies Committee. Mice were maintained on a 12 hour light/dark cycle at 18-26°C in an AAALAC accredited facility. Both male and female littermates ranging in age from 3-24 weeks were used for experiments. TAM was given to *Umod* ^DEL/+^;*Manf* ^fl/fl^ and *Umod* ^DEL/+^;*Umod* ^CE/+^;*Manf* ^fl/fl^ mice, as well as their WT control littermates *Umod* ^+/+^;*Manf* ^fl/fl^ and *Umod* ^+/+^;*Umod* ^CE/+^;*Manf* ^fl/fl^ at 5 weeks of age, 3 mg per oral gavage on each mouse, every other day for three doses. Doxycyline food (1500 ppm, irradiated) was given to *Umod* ^DEL/+^;P8TA and *Umod* ^DEL/+^;MANF/P8TA mice, as well as their WT control littermates *Umod* ^+/+^;P8TA and *Umod* ^+/+^;MANF/P8TA mice at 4 weeks of age from 4-20 weeks.

##### Generation of UMOD Y178-R186 deletion mutant mice

*Umod* ^DEL/+^ mice were generated by Washington University Transgenic Vectors Core. CRISPR gRNAs for *in vitro* testing were identified using CRISPOR (http://crispor.tefor.net/) and synthesized as gBlocks (IDT). *In vitro* target specific gRNA cleavage activity was validated by transfecting N2A cells with PCR amplified gRNA gblock and Cas9 plasmid DNA (px330, Addgene) using ROCHE Xtremegene HP. Cell pools were harvested 48 hours later for genomic DNA prep, followed by Sanger sequencing of PCR products spanning the gRNA/Cas9 cleavage site and TIDE analysis (https://tide.nki.nl/) of sequence trace files. CRISPR sgRNA and Cas9 protein for injection were purchased from IDT and complexed to generate the ribonucleoprotein (RNP) for injection along with a 200 nucleotide ssODN donor DNA (IDT). Injection concentrations were: 50 ng/μl Cas9, 20 ng/μl gRNA and 20 ng/μl ssODN. C57BL/6J F1 mice at 3-4 weeks of age (The Jackson Laboratory, Bar Harbor, ME) were superovulated by intraperitoneal injection of 5 IU pregnant mare serum gonadotropin (SIGMA), followed 48 hours later by intraperitoneal injection of 5 IU human chorionic gonadotropin (SIGMA). Mouse zygotes were obtained by mating C57BL/6J males with superovulated C57BL/6J females at a 1:1 ratio. One-cell fertilized embryos were injected into the pronucleus and cytoplasm of each zygote. Founder genotyping was done by deep sequencing (MiSeq, Ilumina). Mosaic founders were crossed to C57BL/6J (The Jackson Laboratory, Bar Harbor, ME) to generate heterozygous F1 offspring. F1 offspring was deep sequenced to confirm correctly targeted alleles. gRNA sequence: 5’-GCTGCGCCAGTACTCAGTCA-3’. ssODN sequence: 5’- GCACACAGGTCTCAGCCATGCGAACGCCACCCTGGCCTGTGAACCGGTACCAGCCG TGCAGACCCGCGTCACAGGAGTAGCCCACACCATACTCTGTGCTTGTATTGCAGGGGTCTTGACACACCAGCTTTCCATCCGGGCCCTGGGGCAAGCAGTCCAGTCCTGGCTCACAGGAGCCTGGGGAGCACTCACAGTACCAA-3’.

##### Generation of inducible TET-MANF Tg mice

TET-MANF mice were generated by Dr. Maria Lindahl. In brief, to produce these transgenic mice, the full-length cDNA of human MANF was inserted into a *Tet-op-mp1*vector ^48^ (a kind gift of Prof. Harold E. Varmus, Weill Cornell Medicine, NY and Prof. Katerina Politi, New Haven, CT) between a seven direct repeats of the tet-operator sequence (Tet-O_7_) and the mouse protamine 1 poly(A) and intron sequence. The transgenic construct was injected into the pronucleus of fertilized egg cells of FVB/N strain at the Gene Modified Mouse Unit of Laboratory Animal Center, University of Helsinki. Four positive TET-MANF transgenic mouse founders were obtained and transgene copy numbers were analyzed by Southern blotting and quantitative RT-PCR.

*Umod* ^C147W/+^ mice on the C57BL/6 background were described and published previously ^3^.

*Manf* ^fl/fl^ on the C57BL/6 background was generated by Dr. Maria Lindahl and published previously ^18^.

*Umod* ^IRES CRE-ERT^^2^ on the C57BL/6 background was generated by Dr. Andrew P. McMahon in the University of South California and purchased from The Jackson Laboratory (Stock No: 030601). In the CRISPR/Cas9 generated *Umod* ^IRES CRE-ERT^^2^ mice, IRES-CRE-ERT2 was knocked into the 3’ UTR near the stop codon of the *Umod*. *Umod* ^CE/+^ or *Umod* ^CE/CE^ mice do not exhibit any phenotype abnormality for up to 2 years.

Pax8-rtTA mice on the C57BL/6 background were purchased from The Jackson Laboratory (Stock No: 007176). The hemizygous transgenic mice were intercrossed to generate the homozygous transgenic mice for some breedings by utilizing q-PCR. Forward: 5’- TCGATTCTCATGCCCTTCGC-3’, reverse: 5’-GAGCGAGTTTCCTTGTCGTC-3’

### Construct vectors of GFP-UMOD-WT and GFP-UMOD-H177-R185 del

pLenti-CMV-Blast-DEST was obtained from Addgene (#17451) and the UMOD gene was synthesized by Genscript (Piscataway, NJ) and provided in a cloning vector. pLenti-CMV-Blast- SS-eGFP-UMOD was cloned using overlap PCR and Gateway cloning. Here, the eGFP sequence was amplified with a forward primer containing an attB site, signal sequence, and complementary region of eGFP. A reverse primer contained a region complementary to eGFP and a UMOD overhang. The UMOD gene was amplified with a forward primer containing a region complementary to UMOD and an eGFP overhang. The reverse primer contained an attB site and complementary region to UMOD. The eGFP and UMOD genes were then amplified with the respective primers, and then overlap PCR was performed. The amplified product was then inserted into pDONR 221 using a Gateway BP reaction. The donor plasmid, pDONR221- SS-eGFP-UMOD, was then inserted into pLenti-CMV-Blast-DEST via an LR reaction to yield pLenti-CMV-Blast-SS-eGFP-UMOD. The H177-R185 deletion was constructed with a similar approach, with a primer annealing to the respective region to yield the alternative construct.

Cloning was then confirmed via Sanger sequencing. Please note that eGFP was referred to as GFP in the text and figures.

### Lentiviral transduction of HEK 293 cells

Human embryonic kidney (HEK) 293 and HEK 293T cells were cultured at 37°C with 5% CO_2_ in Dulbecco’s modified Eagle medium (DMEM) (Gibco, Carlsbad, USA) containing 2 mmol/L L-glutamine, 100 U/mL penicillin, 100 µg/mL streptomycin, and 10% (vol/vol) fetal bovine serum (Sigma). Lentiviruses were produced by co-transfection of HEK 293T cells with the packaging vector psPAX2 (Addgene #12260), the envelope expression plasmid pMD2.G (Addgene #12259) and lentiviral vector expressing GFP-UMOD-WT or GFP-UMOD-H177- R185 del at a ratio of 3:1:4 using Lipofectamine 3000 Reagent (ThermoFisher Scientific, Waltham, MA) following the manufacturer’s instructions. The medium was changed 8 hours post-transfection. The virus-containing supernatant was harvested and passed through a 0.45-μm filter 72 hours post-transfection. The lentivirus stock expressing GFP-UMOD-WT or GFP- UMOD-H177-R185 del was used for transduction. After HEK 293 cells were infected by lentiviruses, media were changed 24 hours post-transduction and cells were selected with 4 μg/mL Blasticidin (Sigma, CAT 15205) for 3 days after transduction.

### Immunofluorescence staining

For dual staining of MANF or BiP with UMOD in mouse kidneys, 4% paraformaldehyde-fixed paraffin-embedded sections were used. After dewaxing, kidney sections were subjected to antigen retrieval by immersion in 10 mM citric acid buffer (pH 6.0) for 5 minutes at 95°C. Nonspecific binding was blocked by incubating kidney sections with 1% BSA for 30 minutes at room temperature. The slides were stained with a rabbit anti-mouse BiP antibody (Proteintech) or a rabbit anti-mouse (human) MANF antibody (Abnova) together with a rat anti-mouse UMOD antibody (R&D) overnight at 4°C, followed by Alexa 488 or 594-conjugated anti-rabbit or anti-rat secondary antibodies. Slides were analyzed under a fluorescence microscope (Nikon).

For co-staining of MANF with biotinylated LTL in mouse kidneys, paraffin-embedded sections were used. After dewaxing, nonspecific avidin binding was blocked by an avidin/biotin blocking kit (Vector laboratories) before incubating kidney sections with 1% BSA for 30 minutes at room temperature. The slides were stained with a rabbit anti-mouse MANF antibody (Abnova) together with biotinylated LTL overnight at 4°C, followed by an Alexa 488-conjugated anti-rabbit secondary antibody and Alexa 594-conjugated streptavidin, as well as Hoechst 33342. Slides were analyzed under a fluorescence microscope (Nikon).

Formalin-fixed, paraffin-embedded sections of human kidneys underwent the same process of deparaffinization, antigen retrieval and blocking. The slides were incubated with the same rabbit anti-human (mouse) MANF antibody (Abnova), or a rabbit anti-human P62 antibody (Cell Signaling) in combination with a mouse IgG2b antibody against human uromodulin (RayBiotech). A biotinylated anti-mouse antibody was used to amplify fluorescence signals for uromodulin, followed by the corresponding Alexa 488-conjugated anti-rabbit secondary antibody to detect MANF or P62, or by Alexa 594-labelled streptavidin to detect uromodulin. The nuclei were counterstained with Hoechst 33342.

For double staining of FN (Abcam), SMA (Sigma) or F4/80 (Invitrogen) with UMOD in mouse kidneys, frozen sections were employed without antigen retrieval. Ten randomly selected fields (magnification, x400) in each kidney were evaluated, and all images were captured by Olympus DP72 Capture Interface software. Ratio of FN, SMA or F4/80 positive areas to whole areas in each field was calculated in percentages by Image J (NIH) software. TUNEL analysis was performed with the In Situ Cell Death Detection Kit, Fluorescein (Ver16.0, Roche, Mannheim, Germany) per the instructions.

For immunocytochemistry, the stably transduced HEK 293 cells were seeded on coverslips for grow for 48 hours. The cells were fixed in 4% paraformaldehyde for 10 minutes, permeabilized in 0.1% triton for 10 minutes, and blocked with 1% BSA for 1 hour at room temperature, followed by incubation with mouse anti-calnexin antibody overnight at 4°C. The coverslips were washed with PBS and incubated with Alexa 594-conjugated anti-mouse secondary antibody for 1 hour at 37°C. The nuclei were counterstained with Hoechst 33342.

### Light microscopy, electron microscopy and confocal microscopy

For light microscopy, mouse kidneys were fixed in 4% PFA, dehydrated through graded ethanols, embedded in paraffin, sectioned at 4 μm, and stained with Masson’s Trichrome or Picrosirius red by the Morphology Core. For TEM, tissues were fixed, embedded in plastic, sectioned, and stained as described previously ^49^. For live cell imaging, Images were captured using a Nikon Eclipse Ti-E inverted confocal microscope equipped with 10× Plan Fluor (0.30 NA), 20× Plan Apo air (0.75 NA), 60× Plan Fluor oil immersion (1.4 NA), or 100× Plan Fluor oil immersion (1.45 NA) objectives (Nikon). Images were processed and analyzed using Elements AR 5.21 (Nikon).

Fibrosis severity, as quantified by Masson’s Trichrome, was scored in the kidney corticomedullary junction area at ×400 magnification using a counting grid with 140 intersections. The number of grid intersections overlying trichrome-positive (blue) interstitial areas was counted and expressed as a percentage of all grid intersections. For this calculation, intersections that were in tubular lumen and glomeruli were subtracted from the total number of grid intersections ^50^.

### Purification of mouse primary TAL epithelial cells

Mouse UMOD-producing epithelial cells were purified by preparing single cell digests from mouse kidneys of the different genotypes, as described previously ^9^. Briefly, kidneys were decapsulated, minced, and then incubated in a shaking water bath (180 rpm) at 37°C for 50 minutes with Type 1 Collagenase (2 mg/ml; Sigma-Aldrich) and DNase (100 U/ml; Sigma-Aldrich) in serum-free DMEM/F12 (Sigma-Aldrich) with vortexing every 15 minutes. After incubation, enzymatic digestion was halted by addition of 1 volume of DMEM/F12 with 10% serum. The tissue digest was mixed thoroughly and filtered (40 μM) to get the single cell suspension. UMOD producing cells were separated from the single cell suspension using the Dynabead magnetic bead conjugation system, whereby biotinylated, sheep anti-UMOD monoclonal antibody (R&D) was incubated with Dynabeads Biotin Binder (Thermo Fisher, Waltham, MA, CAT 11047). For each kidney isolate, 100 μL beads were conjugated with 4 μg anti-UMOD antibody at 4°C for 45 minutes. Beads were then washed twice with 1% BSA/PBS buffer, and incubated with the single cell suspension in a final volume of 1 ml at 4°C for 45 minutes. Beads and UMOD-positive cells were then trapped against a magnet, and residual UMOD-negative cells were washed away. UMOD-positive fractions were divided for RNA and protein analysis.

### RNA sequencing and analysis

Library preparation was performed with 500 ng of total RNA where the ribosomal RNA was removed by an RNase-H method using RiboErase kits (Kapa Biosystems), fragmented in reverse transcriptase buffer and heated to 94°C for 8 minutes, and then reverse transcribed to yield cDNA using SuperScript III RT enzyme (Life Technologies, per manufacturer’s instructions) and random hexamers. A second strand reaction was performed to yield ds-cDNA. cDNA was then blunt ended, had an A base added to the 3’ ends, and then had Illumina sequencing adapters ligated to the ends. Ligated fragments were then amplified for 12-15 cycles using primers incorporating unique dual index tags, and the indexed fragments for each sample were then pooled in an equimolar ratio and sequenced on an Illumina NovaSeq-6000 using paired end reads extending 150 bases.

Basecalls and demultiplexing were performed with Illumina’s bcl2fastq software and a custom python demultiplexing program with a maximum of one mismatch in the indexing read. RNA-seq reads were then aligned to the Ensembl release 76 primary assembly with STAR version 2.5.1a ^51^. Gene counts were derived from the number of uniquely aligned unambiguous reads by Subread:featureCount version 1.4.6-p5 ^52^. All gene counts were then imported into the R/Bioconductor package EdgeR ^53^ and TMM normalization size factors were calculated to adjust for samples for differences in library size.

Ribosomal genes and genes not expressed in at least 5 samples greater than one count-per-million were excluded from further analysis. The TMM size factors and the matrix of counts were then imported into the R/Bioconductor package Limma ^54^. Weighted likelihoods based on the observed mean-variance relationship of every gene and sample were then calculated for all samples with the voomWithQualityWeights ^55^. Differential expression analysis was then performed to analyze for differences between conditions, and the results were filtered for only those genes with Benjamini-Hochberg false-discovery rate (FDR) adjusted p-values less than or equal to 0.05.

For each contrast extracted with Limma, global perturbations in known Gene Ontology (GO) terms and KEGG pathways were detected using the R/Bioconductor package GAGE ^56^ to test for changes in expression of the log2 fold-changes reported by Limma in each term versus the background log2 fold-changes of all genes found outside the respective term. GO terms and KEGG pathways with Benjamini-Hochberg adjusted p-values less than 0.05 were considered statistically significant.

### Western blot analysis

The isolated UMOD-positive or -negative epithelium and stable HEK 293 cells were lysed by RIPA buffer (Cell signaling CAT9806) containing protease and phosphatase inhibitor cocktails (Roche, Indianapolis, Indiana). The mouse kidneys were extracted using the same lysis buffer with the protease and phosphatase inhibitors and homogenized by sonication. The protein concentrations of cell and kidney lysates were determined by Bio-Rad protein assay (Hercules, CA) using BSA as a standard. Urine volume from individual mouse was normalized to urine creatinine excretion. Denatured proteins were separated on SDS polyacrylamide gel electrophoresis (SDS-PAGE) and then transferred to PVDF membranes. Proteins were not heat denatured for detection of mitochondrial OXPHOS proteins. Blots were blocked with 5% non-fat milk for 1 hour at room temperature, and then incubated overnight with primary antibodies at 4°C. The membranes were washed with Tris-buffered saline/Tween buffer and incubated with the appropriate HRP–conjugated secondary antibodies. The proteins were then visualized in an x-ray developer using ECL plus detection reagents (GE, Pittsburgh, PA) or Chemidoc MP imaging instrument (Biorad). To ensure equal protein loading, the same blot was stripped with stripping buffer (25 mM glycine + 1% SDS, pH = 2.0) and then incubated with a HRP-conjugated anti-mouse β-actin antibody (Sigma) or anti-mouse GAPDH antibody (Cell Signaling, Danvers, MA). Relative intensities of protein bands were quantified using Image J (NIH) analysis software.

### mRNA quantification by Real-Time PCR

Total RNA from primary TALs, stably transduced HEK 293 cells or whole kidneys was extracted using the RNeasy kit (Qiagen, Valencia, CA) with subsequent DNase I treatment. Cellular or kidney RNA was then reverse-transcribed using an RT-PCR kit (Superscript III; Invitrogen). Gene expression was evaluated by quantitative real-time PCR. One µl of cDNA was added to SYBR Green PCR Master Mix (Qiagen) and subjected to PCR amplification (one cycle at 95°C for 20 seconds, 40 cycles at 95°C for 1 second, and 60°C for 20 seconds) in a QuantStudio ^TM^ 6 Flex Fast Real-Time PCR System (Life Technologies, Grand Island, NY). Q- PCR was conducted in triplicate for each sample.

### Preclinical PET/ CT imaging

Radiolabeled ^68^Ga-Galuminox was synthesized using a procedure described previously ^30^. Imaging studies were performed in *Umod ^+/+^* and *Umod* ^DEL/+^ mice at 16 weeks. Both sexes were used. For these studies, mice were anesthetized with isoflurane (2%) via an induction chamber and maintained with a nose cone. After anesthetization, the mice were secured in a supine position and placed in an acrylic imaging tray. Following intravenous tail-vein administration of ^68^Ga-Galuminox (100 μL; 100 μCi; 2% ethanol in saline, 3.7 MBq), static preclinical PET scans were performed over 30-50 minutes, using Inveon PET/CT scanner (Siemens Medical Solutions). PET data were stored in list mode, and reconstruction was performed using a three-dimensional ordered-subset expectation maximization (3D-OSEM) method with detector efficiency, decay, dead time, attenuation, and scatter corrections applied. For anatomical visualization, PET images were also coregistered with CT images from an Inveon PET/CT scanner. Regions of interests were drawn over the kidney, and standard uptake values (SUVs) were calculated as the mean radioactivity per injected dose per animal weight.

### Biodistribution studies

Following micro PET/CT imaging, mice (n=8) were euthanized by cervical dislocation. Blood samples were obtained by cardiac puncture, kidneys harvested rapidly, and all tissue samples analyzed for γ-activity using Beckman Gamma 8000 counter. All samples were decay-corrected to the time, the γ-counter was started. Standard samples were counted with the kidneys for each animal and represented 1% of the injected dose. An additional dose was diluted into milliQ water (100 mL) and aliquots (1 mL) were counted with each mouse. Data were quantified as the percentage injected dose (% ID) per gram of tissue (tissue kBq (injected kBq)^-1^ (g tissue)^-1^ x 100).

### Preparation of permeabilized UMOD^+^ and UMOD^-^ cells and high-resolution respirometry

Isolated UMOD-positive (TAL) and UMOD-negative cells were resuspended in cold mitochondrial respiration solution (MIR05, 0.5 mM EGTA, 3 mM MgCl_2_, 60 mM K-lactobionate, 20 mM taurin, 10 mM KH_2_PO4, 20 mM HEPES, 110 mM sucrose and 1 g/L BSA, pH 7.1) and placed in the Oxygraph 2K (OROBOROS Instruments, Innsbruck, Austria) for high resolution-respirometry measurements ^57–59^. To facilitate uptake of respiratory substrates, cellular plasma membranes were permeabilized in the chamber with digitonin (10 ug/mL) for 20 minutes. Following permeabilization, routine oxygen consumption was measured by the sequential addition of the following substrates: malate (0.5 mM), glutamate (10 mM) and pyruvate (5 mM) to assess complex I-mediated LEAK respiration. Adenosine diphosphate (ADP, 5 mM) to assess maximal complex I respiration followed by succinate (10 mM) to measure OXPHOS (complex I and II-mediated respiration). The uncoupling agent FCCP (Carbonyl cyanide p-trifluoro-methoxyphenyl hydrazone, 0.5 µM, titrated 3X) was then added to determine maximal electron transport system (ETS) capacity. A period of stabilization followed the addition of each substrate. Following assay completion, cells were recovered from the instrument, assayed for total protein content, and then results were normalized to protein amount and analyzed using the DatLab Software (OROBOROS Instruments, Innsbruck, Austria).

### Mitochondrial fractionation

Stably transduced WT and DEL cells were washed with cold PBS first. Mitochondrial and cytosol fractions were isolated from cells using Mitochondrial Isolation Kit for Cultured Cells (Thermo Fisher Scientific) according to the manufacturer’s protocols.

### mtDNA release assay

DNA was isolated from 200 μL of the cytosolic fraction using a DNeasy Blood & Tissue Kit (Qiagen). Quantitative PCR was employed to measure mtDNA using Applied Biosystems SYBR Green Master Mixes (Agilent Technologies) in a QuantStudio ^TM^ 6 Flex Fast Real-Time PCR System. The copy number of mtDNA encoding cytochrome c oxidase I was normalized to nuclear DNA encoding 18S ribosomal RNA. The following primers were used: cytochrome c oxidase I (F: 5′- GAGCCTCCGTAGACCTAACC-3′ and R: 5′- TGAGGTTGCGGTCTGTTAGT-3′) and 18S rRNA (F: 5′- CTCAACACGGGAAACCTCAC-3′ and R: 5′- CGCTCCACCAACTAAGAACG-3′).

### BUN measurement

BUN was measured by using a QuantiChrom^TM^ urea assay kit (DIUR-100) (BioAssay Systems, Hayward, CA).

### Urinalysis

Mouse urines were collected by manual restraint. The mouse urines were centrifuged at 1800g for 10 minutes to remove debris before being frozen at -70°C. Urinary Cr concentration was quantified by a QuantiChrom^TM^ creatinine assay kit (DICT-500) (BioAssay Systems).

### Statistics

Statistical analyses were performed using GraphPad Prism software. Data were expressed as mean ± SD or plots. A 2-tailed Student’s t test was used to compare two groups. One-way ANOVA with post-hoc Tukey test was used to compare multiple groups. *P*<0.05 was considered significant.

### Study approval

All human paraffin-embedded slides from ADTKD patients were obtained from the Wake Forest Cohort under the protocol approved by the Institutional Review Board of Wake Forest School of Medicine.

## Acknowledgments

We thank the Diabetes Research Center Transgenic & ES Cell Core for generating the *Umod* ^DEL/+^ mice (supported by NIH P30 P30DK020579) and Musculoskeletal Research Center Morphology Core (supported by NIH P30AR057235) for histology. We thank Rashmi Nanjundappa, Tao Cheng and Moe Mahjoub for help with confocal live cell imaging. Mice were housed in the Washington University Mouse Genetics Core.

## Funding

E.T. is supported by a NCI Cancer Center Support Grant #P30 CA91842 to the Siteman Cancer Center and by ICTS/CTSA Grant# UL1TR002345 from the National Center for Research Resources (NCRR). T.A.P is supported by Cellular and Molecular Biology Core in the Nutrition Obesity Research Center (P30-DK56341). S.K. is supported by the project LTAUSA19068 from the Ministry of Education, Youth and Sports of the Czech Republic and NU21-07-00033 from the Ministry of Health of the Czech Republic. A.J.B. is supported by NIH grant R21DK106584.

M.E.J. is supported by R35GM128772. F.U. is supported by NIH grants R01 DK112921 and UH2 TR002065. V.S. is supported by grants R0l HL111163, R01 HL142297 and NIBIB P41 EB025815. M.L is supported by Juvenile Diabetes Research Foundation 17-2013-410 and 2- SRA-2018-496-A-B, grants from the Finnish Diabetes Research Foundation and Grants 117044 and 333974 from the Academy of Finland. Y.M.C. is supported by NIH grants R01 DK105056A1, R21DK131557A1, R03DK106451 and K08DK089015, the Office of the Assistant Secretary of Defense for Health Affairs through the Peer Reviewed Medical Research Program under Award W81XWH-19-1-0320, George M. O’Brien Kidney Research Core Centers (NU GoKidney, NIH P30 DK114857; UAB/UCSD, NIH P30 DK079337), The Washington University Office of Vice Chancellor for Research (OVCR) Seed Grant, Mallinckrodt Challenge Grant, Washington University Center for Drug Discovery Investigator Matching Micro Grant and Faculty Scholar Award from the Children’s Discovery Institute of Washington University and St. Louis Children’s Hospital. Y.M.C. is a member of Washington University Institute of Clinical and Translational Sciences (UL1 TR000448) and Diabetes Research Center (NIH P30 DK020579). We thank the Slim Health Foundation and the Black-Brogan Foundation for support. We also thank the Preclinical Imaging Facility at Mallinckrodt Institute of Radiology for help in performing PE/CT imaging studies. The Preclinical Imaging Facility is supported by the Siteman Cancer Center Support Grant (P30CA091842).

## Author contributions

YK and CL designed and performed experiments. CJG designed and performed all experiments in stably transduced HEK 293 cells. ET performed RNA-seq data analysis. AP and MEJ designed and performed cloning of GFP-UMOD-WT and GFP-UMOD-H177-R185 del. TAP conducted high resolution respirometry. JS and VS designed and performed PET/CT imaging. YLF and SJP performed experiments. KK, SK and AB coordinated and supplied patient materials. BGJ and JSD provided *Umod* ^C147W^ mice. FU provided advice in conceiving and conducting experiments. ML generated and provided TET-MANF and *Manf ^fl/fl^* mice. YMC conceived, designed and supervised the study, and wrote the manuscript. All authors contributed to the review and approval of the manuscript.

## Competing interests

Y.M.Chen, S.J.Park, Y.Kim, F.Urano are inventors on a patent entitled “Compositions and methods for treating and preventing endoplasmic reticulum (ER) stress-mediated kidney diseases” (US 11,129,871), which was issued by US Patent and Trademark Office in Sep. 2021.

Y.M.Chen and Y.Kim are inventors on a patent entitled “methods of detecting biomarkers of endoplasmic reticulum (ER) stress-associated kidney diseases” (US 10,156,564), which was issued by US Patent and Trademark Office on Dec. 18, 2018.

J. Sivapackiam and V.S. are inventors on a non-provisional patent, entitled “PET tracers for noninvasive imaging of ROS activity” filed by Washington University in St. Louis, St. Louis, MO. Authors (J.S. & V.S.) declare no competing or financial interests.

## Data and materials availability

All data needed to evaluate the conclusions in the paper are present in the paper and/or the Supplementary Materials.

## Supplementary Materials

**Figure S1.**
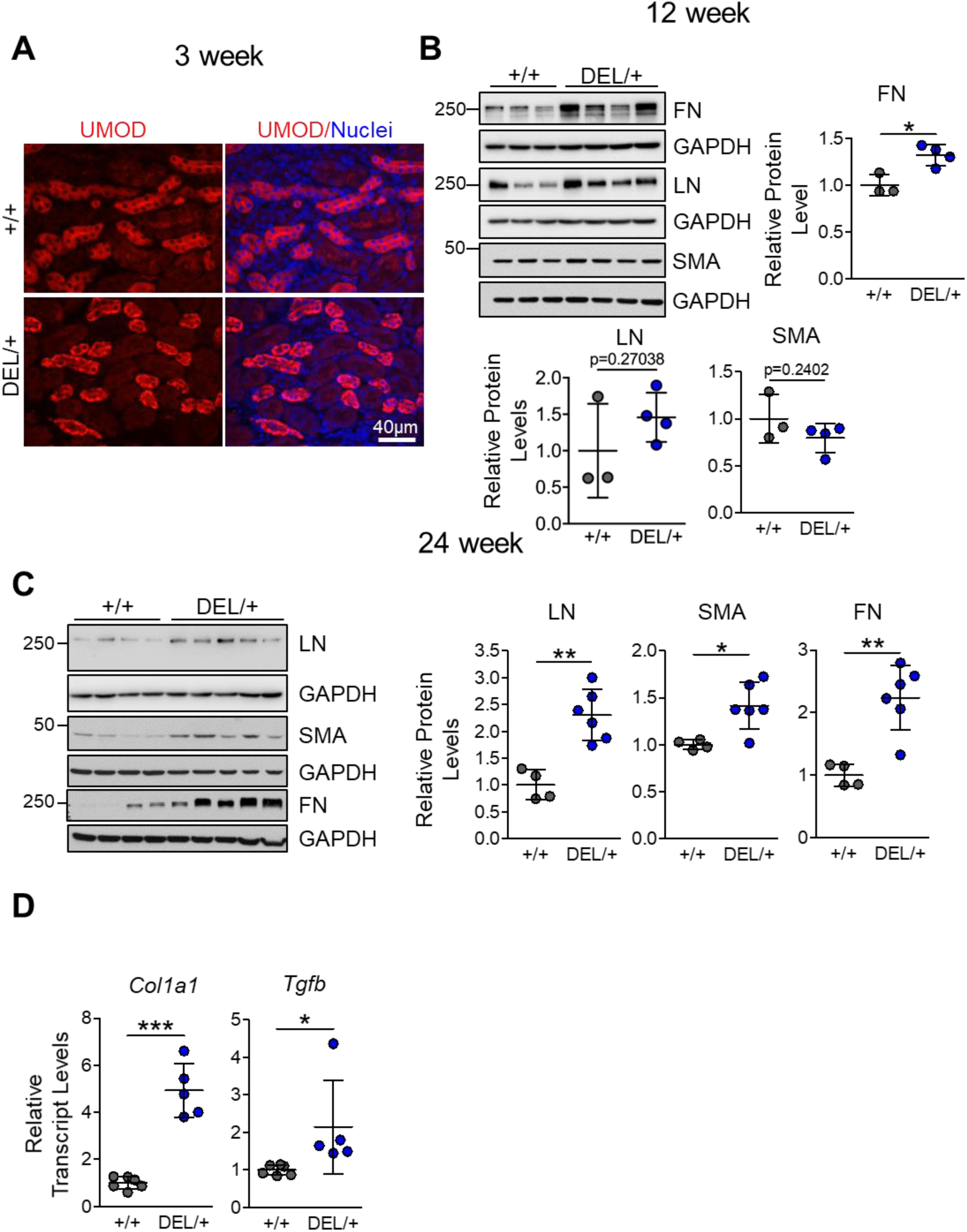
Generation of a mouse model that recapitulates human ADTKD-*UMOD.* **(A)** Representative IF images of paraffin kidney sections stained for UMOD (red) with a nuclear counterstain (Hoechst 33342, blue) at 3 weeks. Scale bar, 40 µm. **(B)** WBs of whole-kidney lysates from *Umod ^+/+^* and *Umod* ^DEL/+^ mice at 12 weeks to detect FN, LN and SMA with densitometry analysis. Mean ± SD (n=3-4 mice/genotype). *p<0.05. **(C)** Representative immunoblots of whole-kidney lysates from *Umod ^+/+^* and *Umod* ^DEL/+^ mice at 24 weeks to detect LN, SMA and FN with densitometry analysis. Mean ± SD (n=4-6 mice/genotype). *p<0.05; **p<0.01. **(D)** Quantitative RT-PCR analysis of relative transcript levels of *Col1a1* and *Tgfb* in whole kidneys from *Umod ^+/+^* and *Umod* ^DEL/+^ mice at 24 weeks. Gene expression was normalized to 18s. Mean ± SD (n=5-6 mice/genotype). *p<0.05; ***p<0.001.

**Figure S2.**
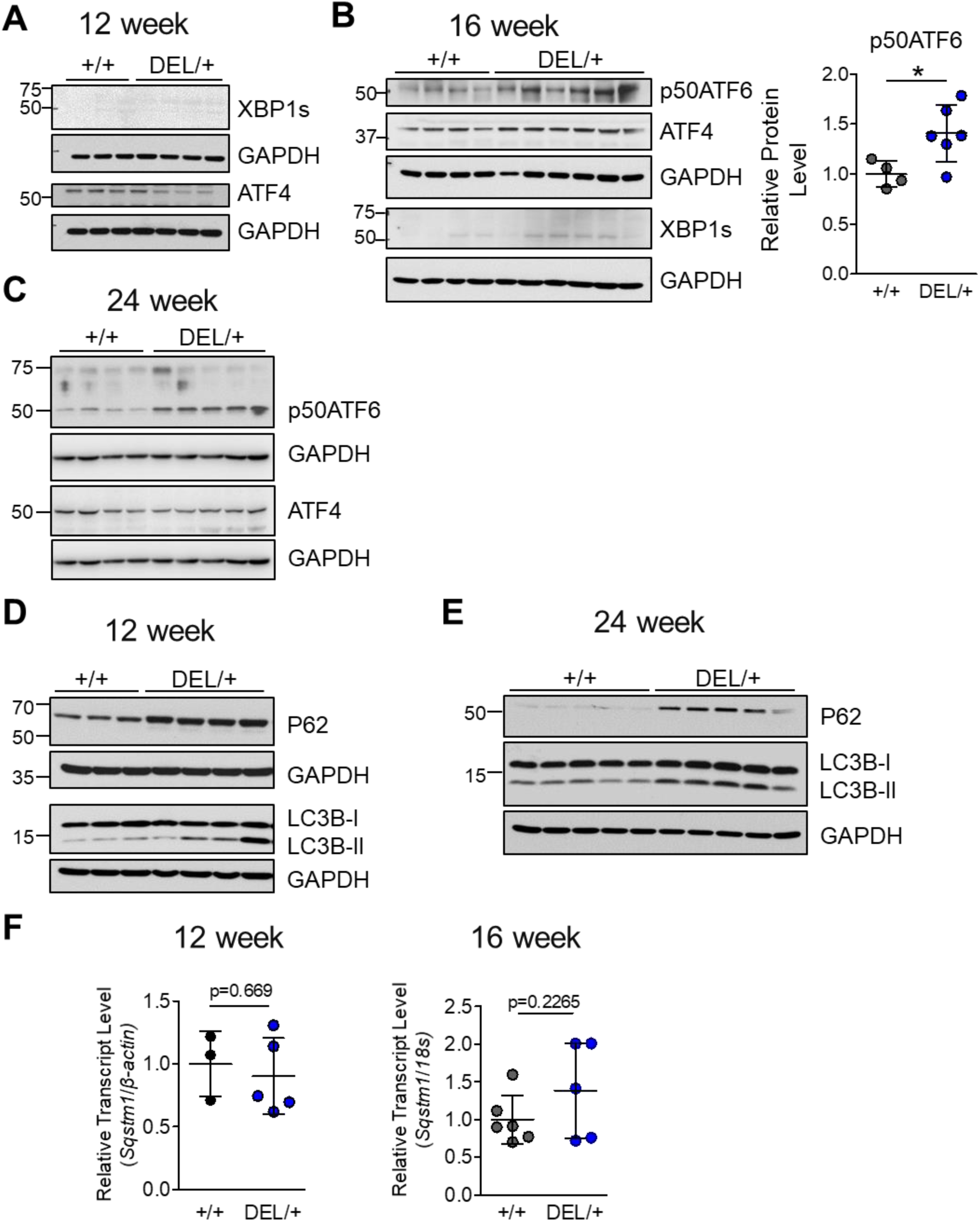
ER stress activation and impaired autophagy in the mouse model of ADTKD-*UMOD*. **(A)** Whole-kidney lysates from *Umod ^+/+^* and *Umod* ^DEL/+^ mice at 12 weeks were examined by WBs for XBP1s and ATF4. n= 3-4 mice/genotype. **(B)** Whole-kidney lysates from *Umod ^+/+^* and *Umod* ^DEL/+^ mice at 16 weeks were examined by WBs for p50ATF6, ATF4 and XBP1s. n=4-6 mice/genotype. For densitometry analysis of p50ATF6, Mean ± SD (n=4-6 mice/genotype). *p<0.05. **(C)** Whole-kidney lysates from *Umod ^+/+^* and *Umod* ^DEL/+^ mice at 24 weeks were examined by WBs for p50ATF6 and ATF4. n=4-5 mice/genotype. **(D-E)** WB of whole-kidney lysates to detect the autophagy mediators P62 and LC3B at 12 weeks, n=3-4 mice/genotype (**D**) and 24 weeks, n=5 mice/genotype (**E**). **(F)** Quantitative PCR of primary TAL cells at 12 and 16 weeks for P62. Mean ± SD (n=3-6 mice/group).

**Figure S3.**
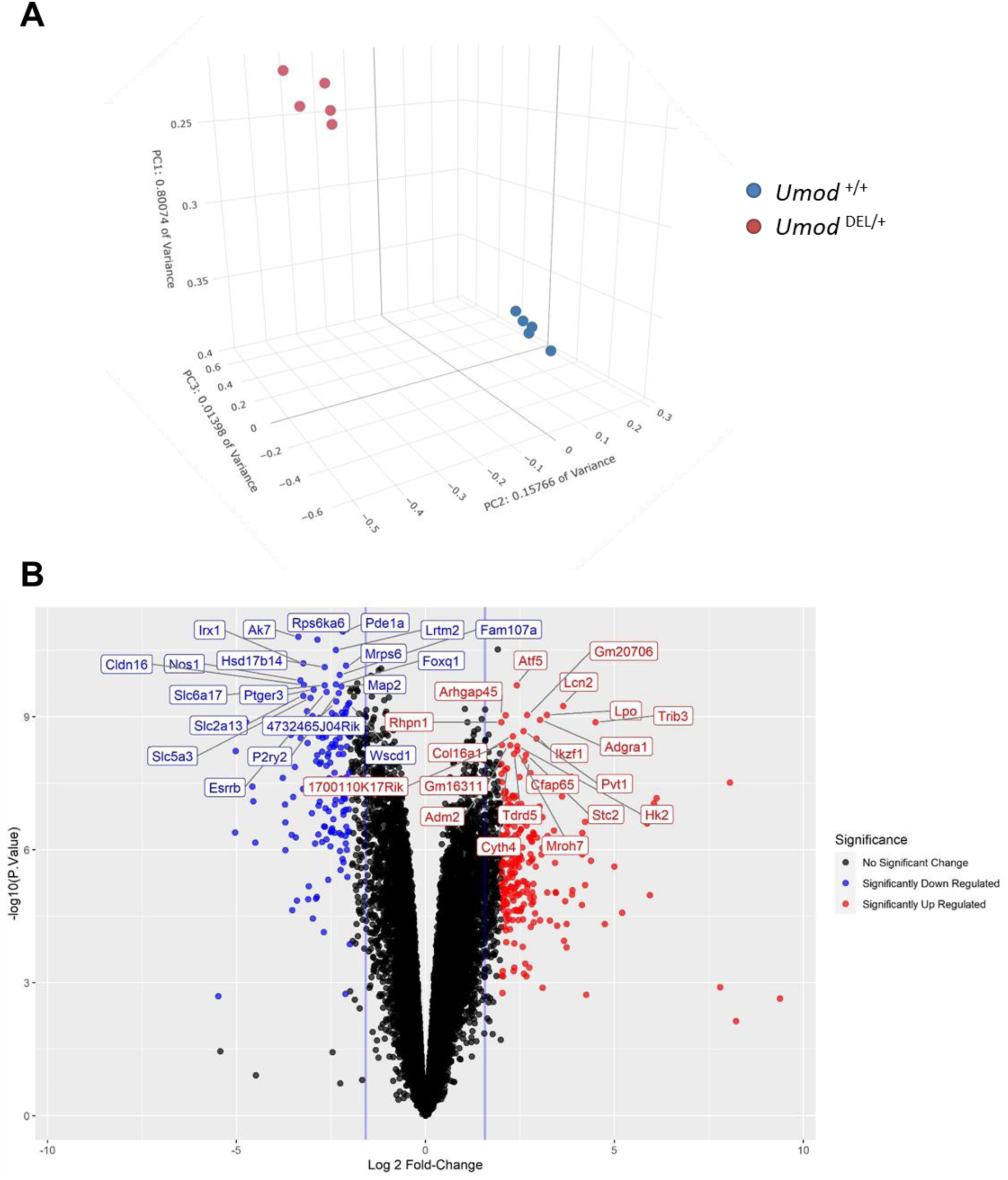
The altered transcriptional profile by the *Umod* mutant allele in primary murine TAL cells at 16 weeks. **(A)** Multidimensional principal component analysis of RNAseq data obtained from *Umod* ^DEL/+^ (n=5) and WT (n=5) TALs at 16 weeks. **(B)** Volcano plot of log2 fold-change and -log10 scale p values, derived by Limma analysis for differential gene expression (Benjamini-Hochberg adjusted P values≤0.05). 9566 were differentially expressed between mutant and WT TAL cells (FDR≤0.05). Out of these transcripts, 244 were up-regulated (red) and 149 were down-regulated (blue) by 4-fold or more. The top 20 most significantly differentiated genes were labeled.

**Figure S4.**
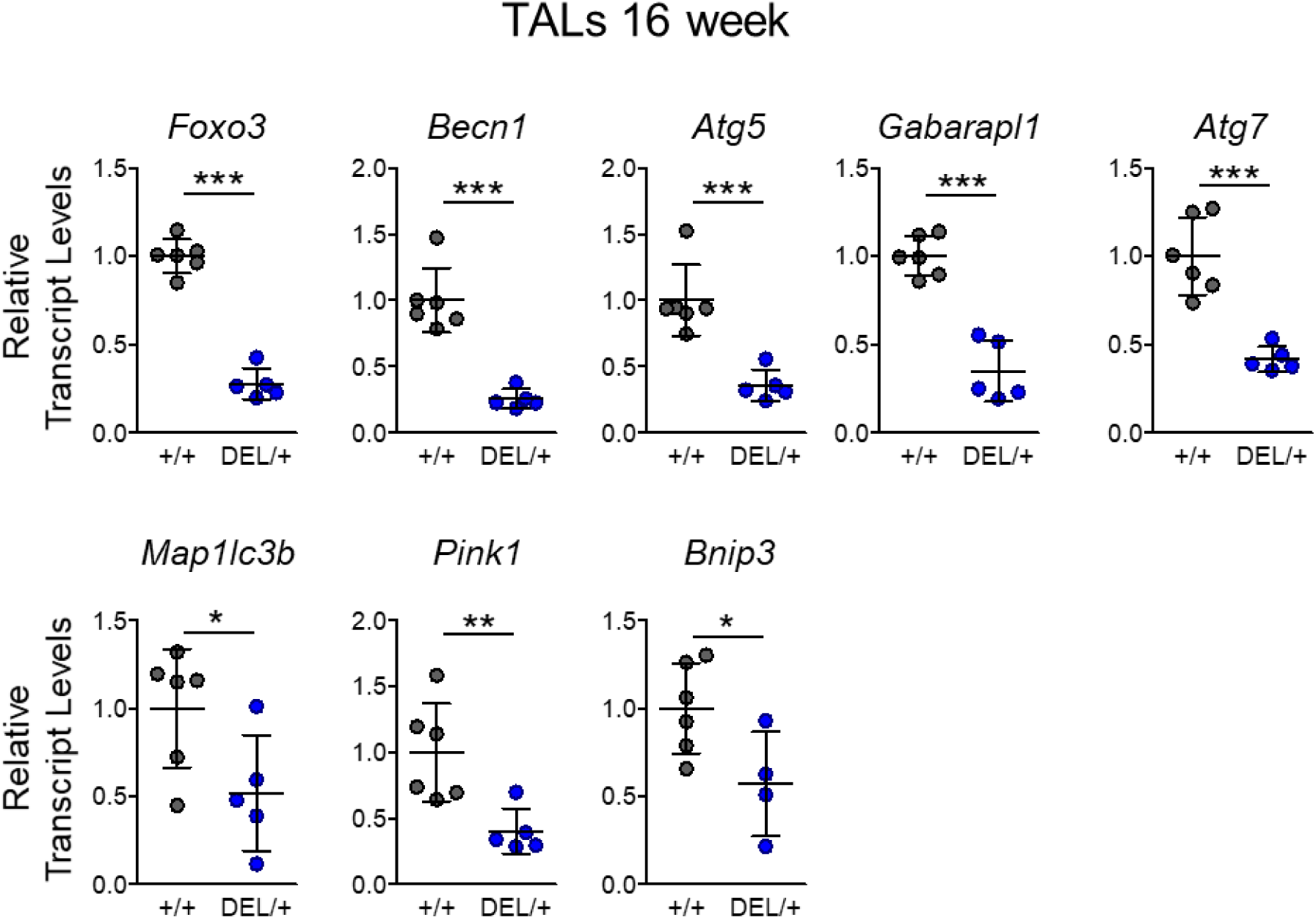
Quantitative PCR validation of RNA-seq results in TAL cells related to autophagy suppression at 16 weeks. Quantitative PCR for a panel of autophagy-related genes of isolated primary TAL cells at 16 weeks. Gene expression was normalized to 18s. Mean ± SD (n=5-6 mice/genotype). *p<0.05; **p<0.01; ***p<0.001.

**Figure S5.**
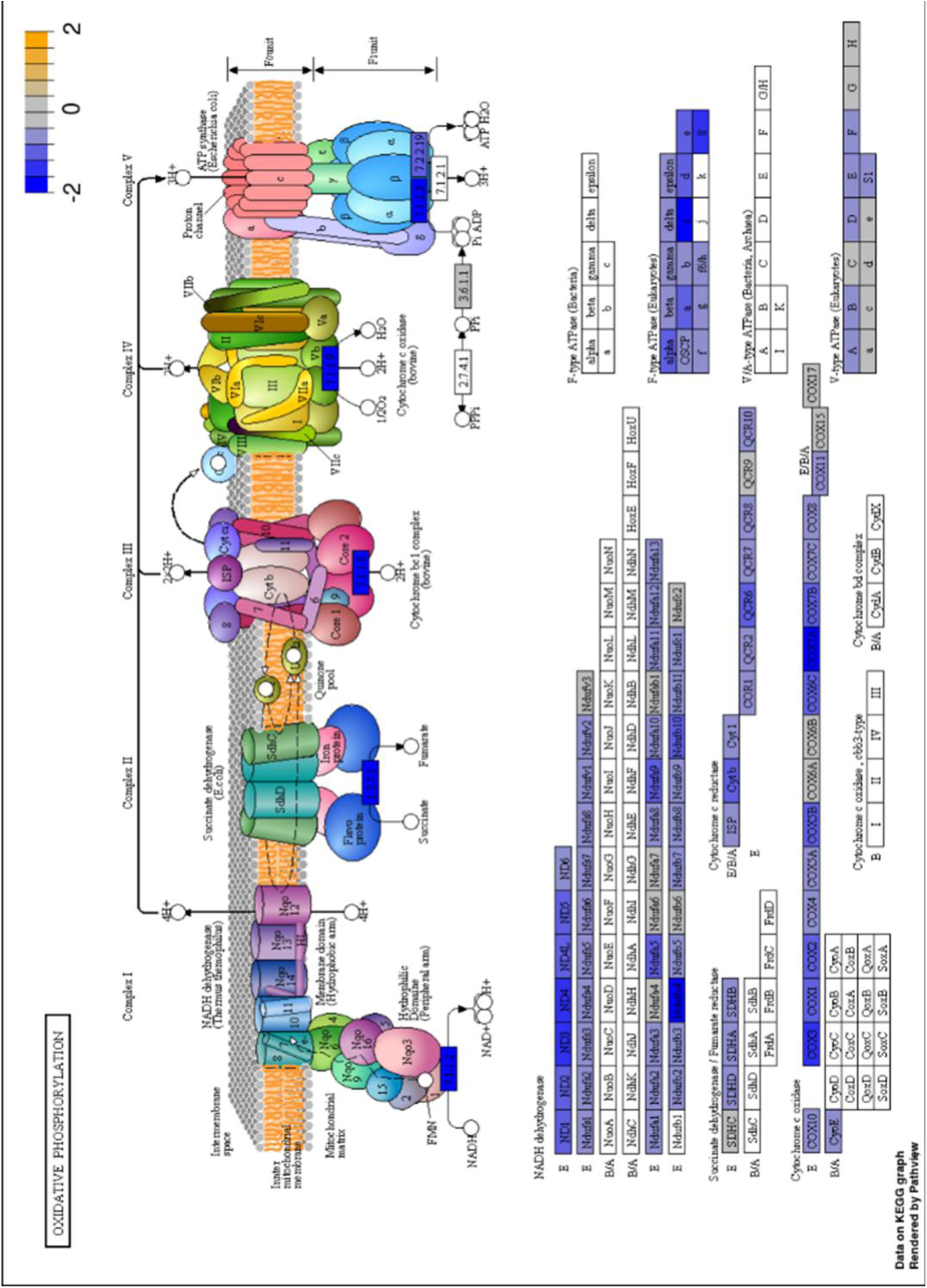
KEGG pathway analysis of mitochondrial respiratory complexes in *Umod* ^DEL/+^ TALs compared with *Umod* ^+/+^ TALs at 16 weeks. The KEGG pathway depicted downregulation of all five ETCs in the mutant TALs at 16 weeks. Color indicates the magnitude of gene expression changes. The most upregulated genes are highlighted in orange, whereas the most downregulated genes are in blue.

**Figure S6.**
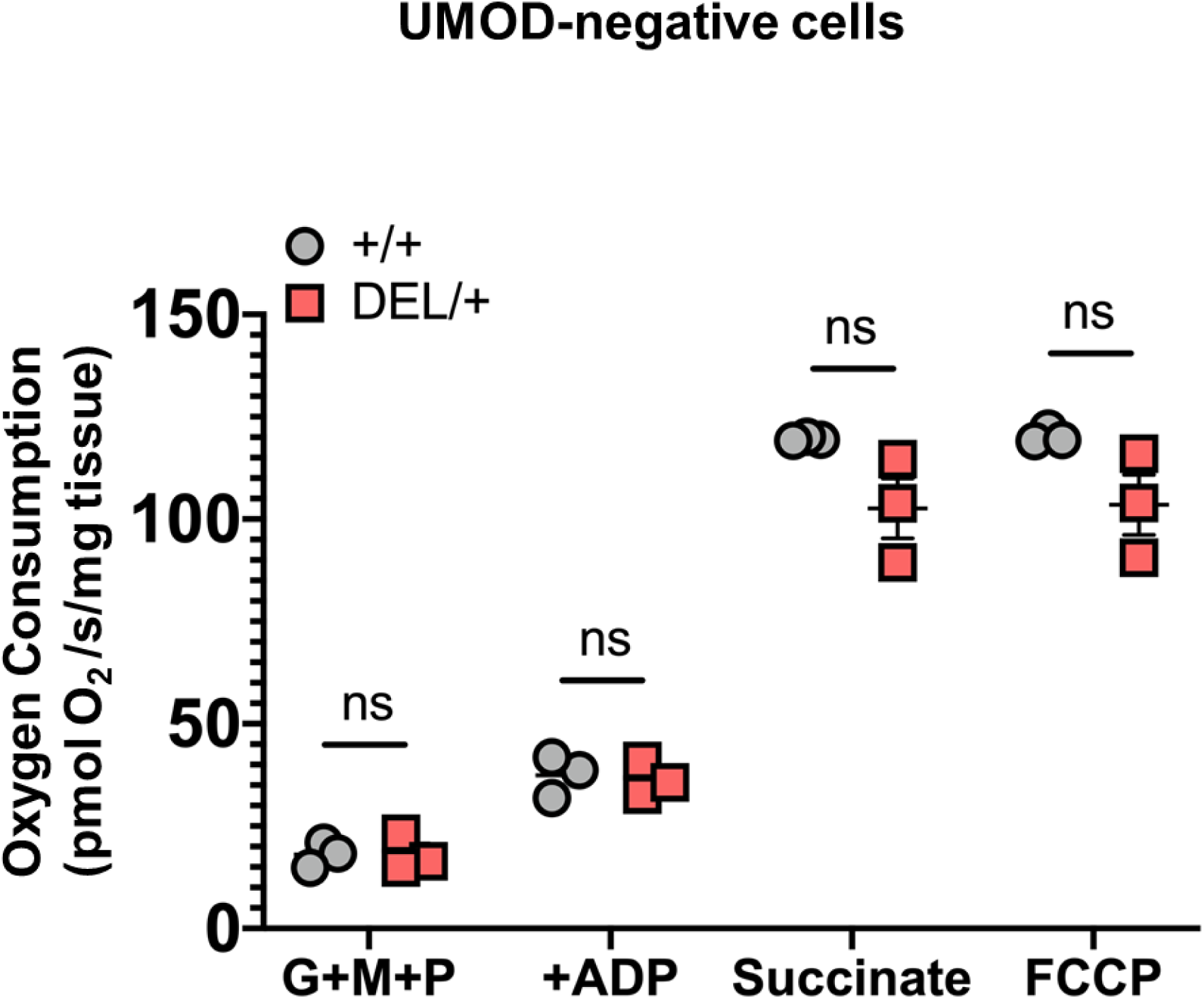
Mitochondrial respiratory function in UMOD^-^ cells isolated from *Umod* ^+/+^ and *Umod* ^DEL/+^ kidneys at 16 weeks. Measurement of mitochondrial respiration using an OROBOROS Oxygraph system in permeabilized UMOD-negative cells from *Umod ^+/+^* and *Umod* ^DEL/+^ kidneys at 16 weeks following sequential additions of glutamate, malate and pyruvate (G+M+P); adenosine diphosphate (ADP); succinate and FCCP. Mean ± SD (n=3 mice/genotype). ns, not significant.

**Figure S7.**
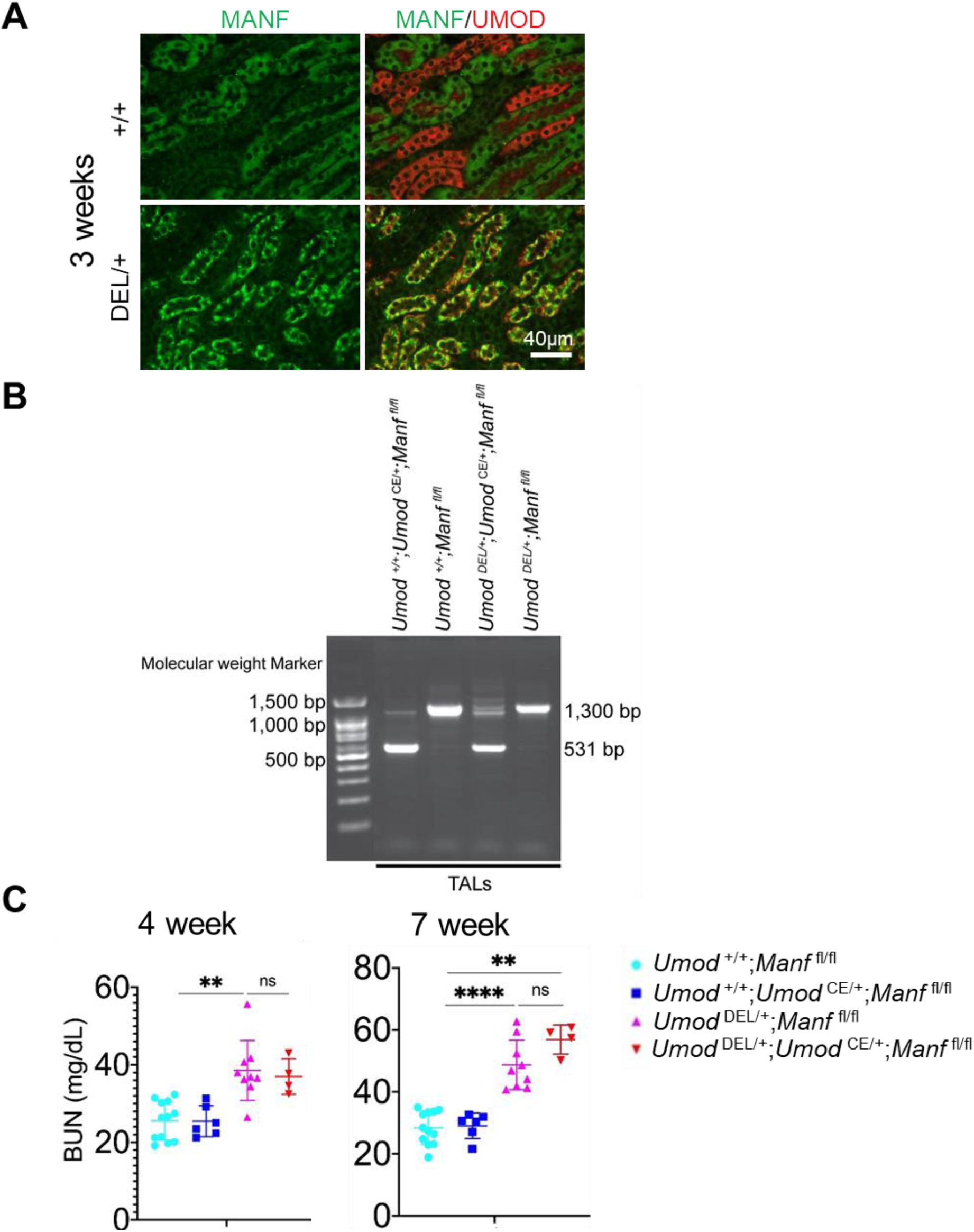
Loss of MANF in TALs deteriorates autophagy suppression and kidney fibrosis in ADTKD. **(A)** Double IF staining for MANF (green) and UMOD (red) on paraffin kidney sections from *Umod ^+/+^* and *Umod* ^DEL/+^ mice at 3 weeks of age. Scale bar, 40 µm. **(B)** PCR results of genomic DNAs from isolated TALs of the indicated genotypes at 12 weeks. All mice were given TAM at 5 weeks of age. Forward primer: 5‘-TGG AGT GAG CAC AAC TCA GG-3’; Reverse primer: 5‘-TGC CATGGT GAT GCT GTA AC-3’. **(C)** BUN measurements at 4 and 7 weeks. Mean ± SD (n=4-12 mice/genotype). ns, not significant; **p<0.01; ****p<0.0001. All mice were given TAM at 5 weeks of age.

**Figure S8.**
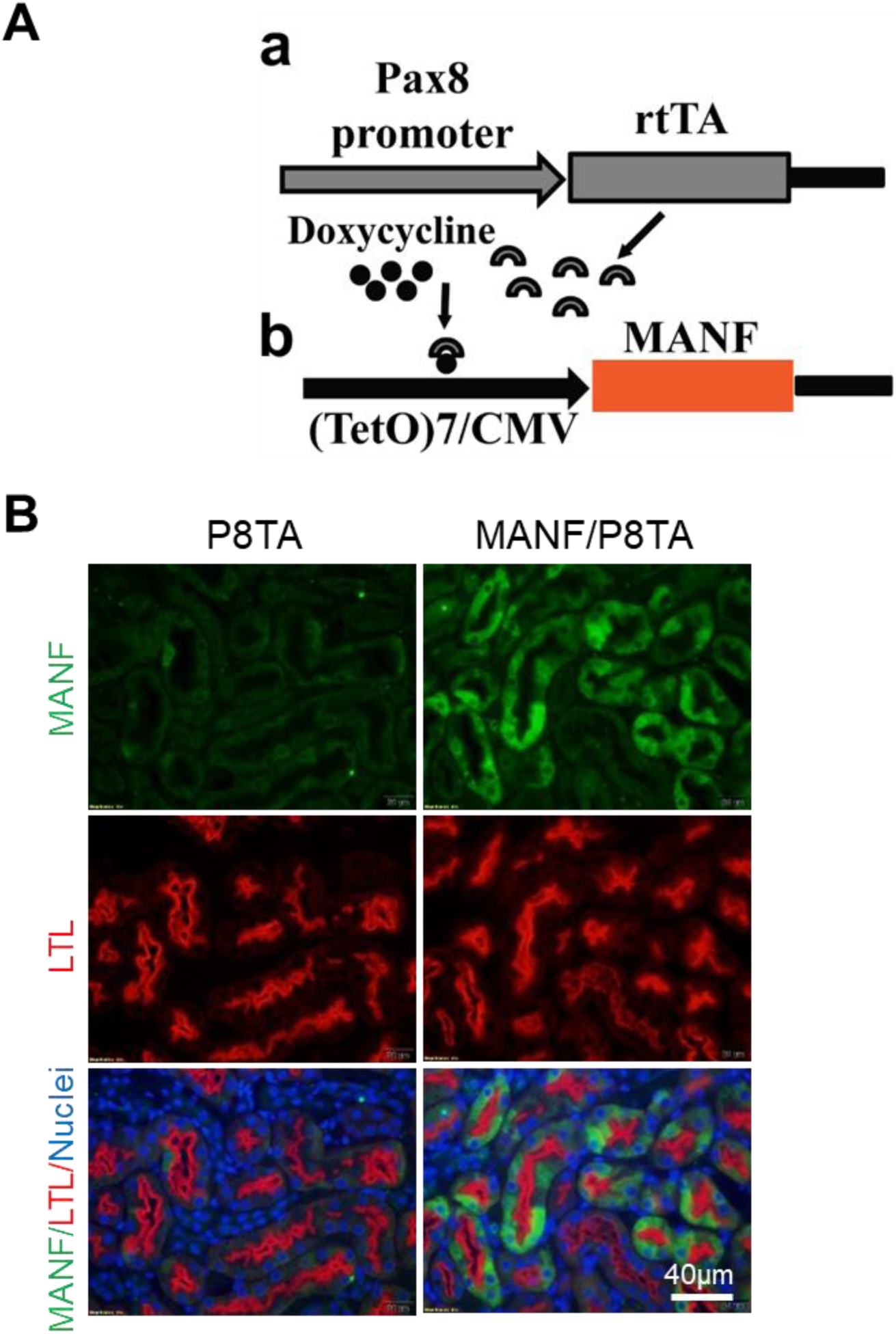
Generation and characterization of inducible, renal tubular MANF transgenic mice. **(A)** Diagram of the two constructs used in the Tg mouse lines for the generation of the Tet-On system. a. The Pax8 promoter directs expression of rtTA in all renal tubular epithelial cells. b. The (TetO)7/CMV-MANF Tg is induced by DOX-bound rtTA. **(B)** IF staining of MANF (green), LTL (red) and Hoechst 33342 (blue) on kidney paraffin sections of the indicated groups after 4 weeks of DOX administration. Scale bar, 40 µm.

**Figure S9.**
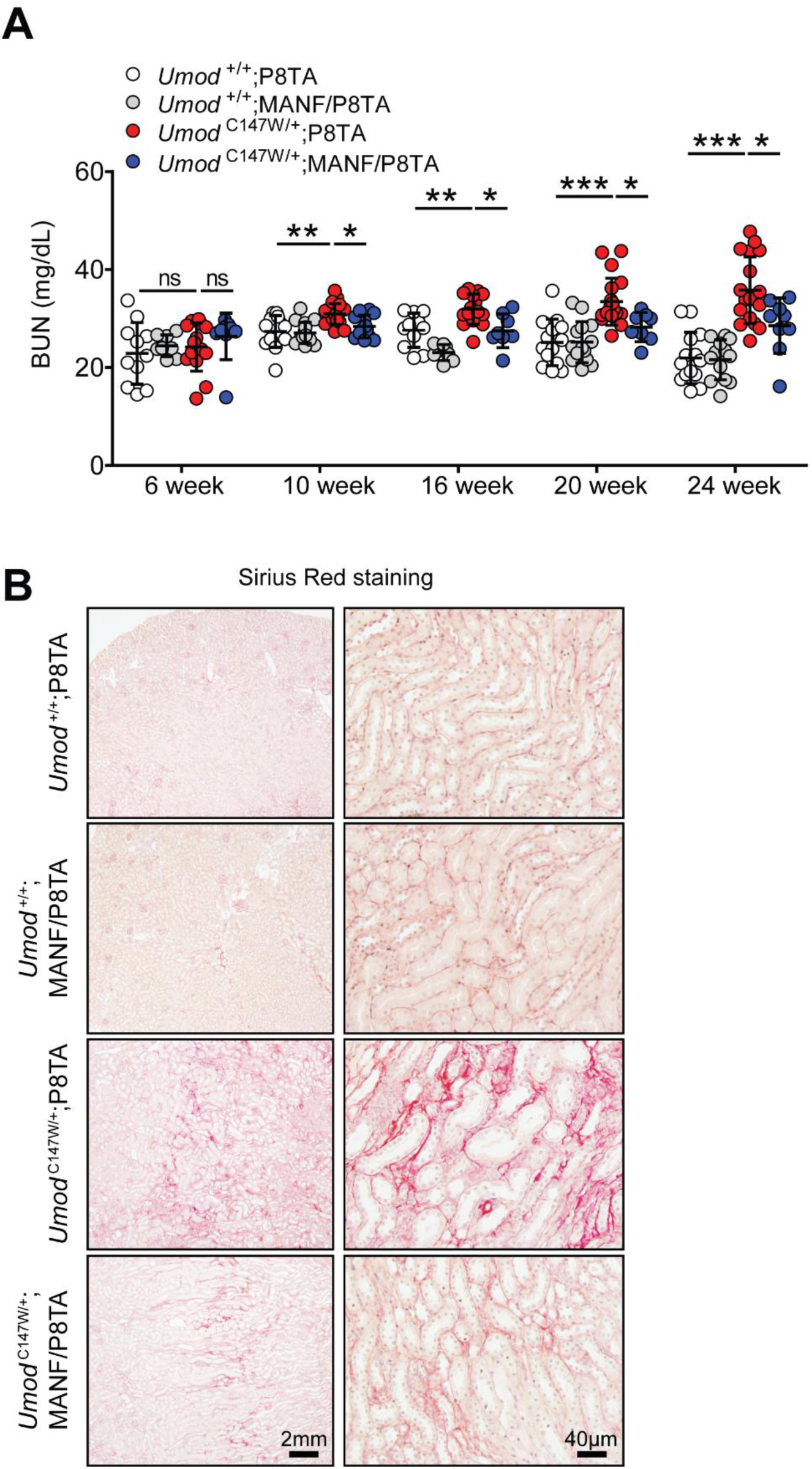
Tubular MANF overexpression inhibits renal fibrosis and protects kidney function in *Umod* ^C147W/+^ mice. **(A)** BUN measurements in the indicated groups over a 24-week period. Mean ± SD (n=8-19 mice/genotype). ns, not significant; *p<0.05; **p<0.01; ***p < 0.001. All mice were given DOX starting from 6 weeks. **(B)** Representative histological images of whole kidney sections stained with Picrosirius red (collagens I/III in red) at 24 weeks. Scale bar, 2 mm or 40 µm.

